# Multiplexed and millimeter-scale fluorescence nanoscopy of cells and tissue sections via prism-illumination and microfluidics-enhanced DNA-PAINT

**DOI:** 10.1101/2022.08.07.503091

**Authors:** Matthew J Rames, John Kenison, Daniel Heineck, Fehmi Civitci, Malwina Szczepaniak, Ting Zheng, Julia Shangguan, Yujia Zhang, Sadik Esener, Xiaolin Nan

## Abstract

Fluorescence nanoscopy has become increasingly powerful for biomedical research, but it has historically afforded a small field-of-view (FOV) around 50 µm x 50 µm at once and more recently up to ~200 µm x 200 µm. Efforts to further increase the FOV in fluorescence nanoscopy have thus far relied on the use of fabricated waveguide substrates, adding cost and sample constraints on the applications. Here we report PRism-Illumination and Microfluidics-Enhanced DNA-PAINT (PRIME-PAINT) for multiplexed fluorescence nanoscopy across millimeter-scale FOVs. Built upon the well-established prism-type total internal reflection microscopy, PRIME-PAINT achieves robust single-molecule localization with up to ~520 µm x 520 µm single FOVs and 25-40 nm lateral resolutions. Through stitching, nanoscopic imaging over mm^2^ sample areas can be completed in as little as 40 minutes per target. An on-stage microfluidics chamber facilitates probe exchange for multiplexing and enhances image quality particularly for formalin-fixed paraffin-embedded (FFPE) tissue sections. We demonstrate the utility of PRIME-PAINT by analyzing ~10^6^ caveolae structures in ~1,000 cells and imaging entire pancreatic cancer lesions from patient tissue biopsies. By imaging from nanometers to millimeters with multiplexity and broad sample compatibility, PRIME-PAINT will be useful for building multiscale, Google-Earth-like views of biological systems.

## Introduction

Advances in superresolution microscopy techniques such as PALM^1,2^, STORM^3^, PAINT (including DNA-PAINT)^4,5^, STED^6^, and their relatives^7,8^ in the recent decades have pushed the effective resolution of optical microscopy to ~20 nm or better, providing unparalleled insights into biological structures and processes at the molecular scale. Until very recently, however, the various fluorescence nanoscopies have afforded rather small FOVs, commonly ~50 µm x 50 µm in the lateral dimensions, addressing only few cells at once. This has limited the utility of fluorescence nanoscopy in studying complex and heterogeneous biological systems, for example multicellular structures in model systems or clinical samples, which can comprise hundreds of cells or more and span millimeters and beyond.

Emerging research has begun to tackle this issue. Huang et al. used a high power illumination scheme to obtain FOVs up to ~200 µm x 200 µm with STORM^9^. In this case, ~6W laser power (at 640 nm) was necessary for efficient fluorophore photoswitching across such large illumination areas. Utilizing a sophisticated line-scanning illumination scheme termed ‘ASTER’, Mau et al. also reached FOVs up to ~200 µm x 200 µm at a moderate laser power (~250 mW) for both STORM and DNA-PAINT^10^. As in most fluorescence nanoscopies, these methods employed the popular, through-objective imaging scheme where the same, high numerical aperture (NA >1.4) objective supports both sample illumination and signal collection. Despite the high signal collection efficiency and resolving power of these objective lenses, the best achievable FOV is typically not more than ~200 µm x 200 µm, beyond which optical aberrations and beam geometry constraints become limiting. At this FOV, tens of (5x5 or more) FOVs need to be stitched to reach the millimeter scale, which will still take hours to days to complete even for a single target.

Even larger FOVs in fluorescence nanoscopies would likely need to involve imaging schemes where separate light paths are used for sample illumination and signal detection. Indeed, several recent reports based on planar waveguides have demonstrated FOVs up to ~0.5 mm x 0.5 mm. These methods employ high-refractive index contrast (HIC) materials such as Ta_2_O_5_ or Si_3_N_4_ ^11–13^ grown on silicon or polymer^14^ deposited on clear glass to create a thin, planar waveguide as the imaging substrate. Excitation light is coupled to the waveguide from the side using specialized optics (e.g. a dedicated objective lens) to generate a continuous, evanescent field to illuminate the sample along the beam path. The fluorescence signal can be collected using an objective of choice, allowing imaging at different magnifications and resolution levels. Typically, by using a high power (e.g. 60x and NA 1.2-1.5) objective, lateral resolutions of ~50 nm could be achieved (e.g. with STORM) over a smaller (~200 µm x 200 µm) FOV^11–13^. Larger FOVs up to ~0.5 x 0.5 mm could be obtained at a lower magnification and NA (e.g. 25x NA 0.8), with lateral resolutions in the 70-200 nm range^13^. The platform is also compatible with other sub-diffractive imaging modalities such as superresolution radial fluctuation (SRRF)^14^.

Despite the impressive FOVs with chip-based nanoscopy, routine use of this platform in most biomedical laboratories can be faced with several hurdles, with the first being the need for using microfabricated HIC waveguides and dedicated alignment optics for light coupling. By design, the input light also needs to be under total-internal-reflection (TIR) throughout the entire light path, which will limit the use under non-strict TIR illumination conditions. The light-coupling optics (e.g. the objective lens) often needs to be mechanically dithered to overcome interference patterns due to repeated reflections of light at the interface^12^. An additional consideration is sample compatibility. While the waveguide substrates have shown good performance on cultured cells and cryo-preserved tissue sections^15^, its applications to formalin-fixed paraffin-embedded (FFPE) tissue sections, the most common clinical sample format, remains to be developed. In previous work, we and others have demonstrated the utility of fluorescence nanoscopy in revealing structural and molecular details such as chromatin organization in early cancer progression^16^ and mitochondria organization^17^ from FFPE sections. Extending fluorescence nanoscopy to FFPE tissue sections with large-field capability in ways that can be seamlessly integrated into current, standard workflows would be of immense value.

To address these needs, we have developed a microscope platform for multiplexed imaging of cells and clinical tissue samples with DNA-PAINT across up to ~0.5 mm x 0.5 mm FOVs in a single acquisition. This platform, which we termed PRism-Illumination and Microfluidics-Enhanced (PRIME-) PAINT, utilizes a prism-based illumination scheme that can be easily implemented on common, inverted fluorescence microscopes with off-the-shelf optical components. PRIME-PAINT affords FOVs equivalent to chip-based nanoscopy but maintains spatial resolutions comparable to that of through-objective systems. Multiplexed, nanoscopic imaging via exchange-PAINT is facilitated using an integrated microfluidic sample chamber, which was critical to high quality imaging of clinical FFPE tissue sections through microfluidics-enhanced DNA-PAINT. We demonstrate multicolor imaging of both cultured cells and clinical, FFPE sections with nanometer (25-40 nm) spatial resolutions. Enabled by the dramatically increased data throughput, we used machine learning-based image segmentation and quantitation of the resulting, multiscale nanoscopic images, offering new insights into nanostructures and molecular interactions across larger cell populations. Collectively, we present PRIME-PAINT as a novel approach to high-resolution and high-throughput spatial mapping of cells and clinical samples across the scales from molecules to multicellular systems.

## Results

### Integrating prism-type illumination and microfluidics with DNA-PAINT

PRIME-PAINT combines a prism-type illumination scheme with an on-stage microfluidic system for programmable fluid exchange. This combination was essential to enabling multiplexed fluorescence nanoscopy via exchange-PAINT over much larger FOVs compared with prior configurations. We chose DNA-PAINT as the primary nanoscopy method in this work because of its convenient multiplexing through exchange-PAINT^5^ and robust multi-FOV stitching without probe loss during imaging^18^. In this work, we specifically used DNA-PAINT-ERS^19,20^, our improved implementation of DNA-PAINT, where the combination of ethylene carbonate (<u>E</u>C, as an imaging buffer additive), repetitive docking sequences, and a <u>s</u>pacer between the affinity agent and the docking strand (DS) oligo enables fast and high-quality DNA-PAINT without needing strong laser excitation. Although not yet tested, the platform should also be compatible with other accelerated implementations of DNA-PAINT^21–23^.

Two TIR configurations, namely prism-illumination^24^ and through-objective^25^, are widely used for single molecule imaging; both are also compatible with highly inclined laminar optical (HiLo) microscopy. Of the two, prism-illumination does not rely on TIR-capable objectives and thus is more flexible on the achievable FOV. Of note, current prism-illumination setups for single molecule imaging still uses high NA (commonly 60x oil, NA=1.4 or above) objectives and deliver FOVs around 50 µm x 50 µm. In our implementation, the laser first passes a pair of beam expanders, one of which installed in reverse, to yield a collimated beam of adjustable size (**Fig. 1A**). A cage-mounted 4π lens pair was placed between a rotatable mirror and the prism such that the FOV center remains static when rotating the mirror to adjust the incident angle of the beam. This setup allowed us to continuously adjust the illumination beam size between tens of µms and millimeters and switch between TIR and HiLo modes without needing to re-center the FOV. It also avoids focusing of high lasers onto sensitive optical surfaces (e.g. the back of an objective lens) and thus can accommodate more incident power if needed (**Fig. 1A**).

**Fig. 1:**
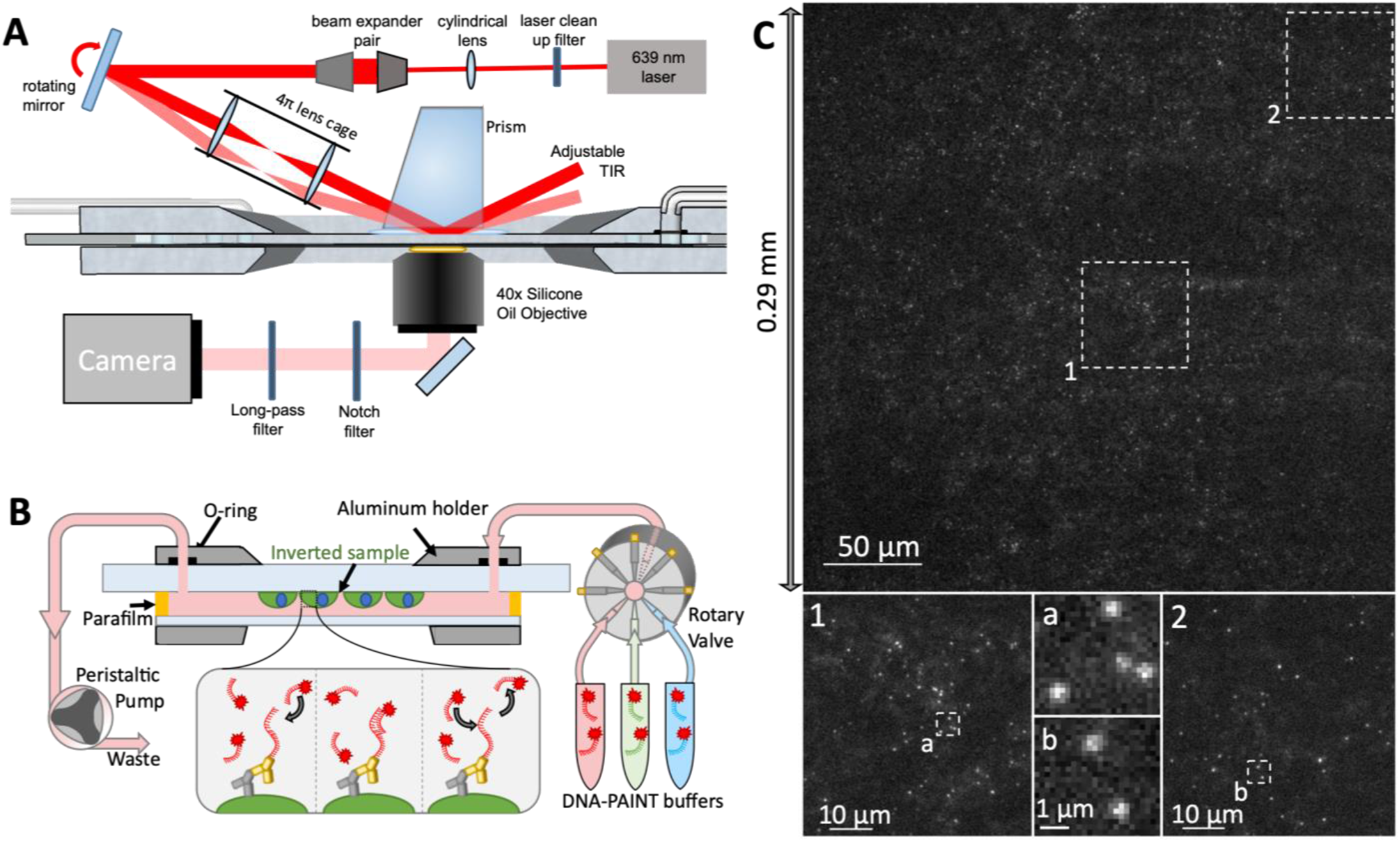
Overview of PRIME-PAINT optics, fluidics, and example SMLM images. (**A**) Schematic of the PRIME-PAINT microscope design and general imaging strategy, with the widened laser beam and controllable TIR angle; (**B**) Representative PRIME-PAINT raw image at 30 ms exposure highlights single-molecule localization quality. Single-molecule events are clearly resolved both in the center on the FOV, and near the corners, exemplifying the 300 µm x 300μm FOV; (**C**) Schematic cross-section of the flow chamber assembly and sample position. Notably, cultured cells/mounted tissue are attached on an inverted sample coverslide, enabling both encapsulation of imaging solutions and buffer exchange for multi-target DNA-PAINT imaging. DNA-PAINT imaging utilizes oligonucleotide-conjugated antibodies (DNA in red, antibody in yellow) and complimentary imaging strand (IS, also shown in red with a fluorophore represented by a red star) to provide transient single-molecule localizations while freely diffusing strands contribute only slightly to background fluorescence.

With sample illumination decoupled from signal detection, we could choose objective lenses best suited for single-molecule detection across large FOVs and at long working distances (**Fig. 1A-B**). We found that, on our setup, a 40x silicone (Sil) immersion objective^26^ (NA 1.25) outperformed the more commonly used 60x oil immersion (NA=1.4) objective. Despite the lower NA, the Sil objective yielded nearly identical single-molecule brightness as the oil objective while affording a lower background and better overall signal-to-noise ratio. Using the Prime 95B sCMOS with matching (60x) magnifications, the image quality started to deteriorate beyond the central ~100 µm x100 µm region with the oil objective but remained uniform across the entire 291 µm x 291 µm FOV with the Sil objective (**Supplementary Fig. S1** and **Fig. 1C**). By switching to another sCMOS, the Kinetix, we achieved an even larger FOV of 521 µm x 521 µm with comparable single molecule image quality (**Supplementary Fig. S2**).

To accommodate this imaging configuration, the samples is sandwiched between a glass slide and a coverglass with the space in between filled with imaging buffer. We designed a microfluidic flow cell with a tear-drop shape for efficient and complete buffer exchange in multiplexed DNA-PAINT (exchange-PAINT)^5^, inspired by previously demonstrated designs^27^. Fire-polished glass slides with low background could be directly used with conventional tissue cell-culture or coated with polyethyleneimine (PEI) for use with FFPE tissue sections (see *Methods*). Stretched and molten parafilm was used as a thin (~35 ± 5 µm) spacer and a robust, hydrophobic seal. An optional, custom holder (see CAD file, **Supplementary data**) helps support and position the sample chamber on the microscope stage with improved mechanical stability. Finally, inlet and outlet fluid ports were connected to a rotary valve and a peristaltic pump, allowing manual or programmable fluid exchange during imaging **(Fig. 1B)**.

### Multiplexed PRIME-PAINT for membrane and cytosolic targets in cultured cells

High-quality single molecule imaging with PRIME-PAINT enabled large-field, nanoscopic imaging of cellular targets in cultured cells with single FOVs of ~0.3 mm x 0.3 mm (**Fig. 2**) or ~0.5 mm x 0.5 mm (**Supplementary Fig. S2-3**). Imaging can be altered between membrane (such as caveolin; **Fig. 2A-D**) and cytosolic (such as mitochondria; **Fig. 2E-H**) targets by switching between TIR and HiLo imaging modes, respectively. The acquisition time for each single FOV was ~10-20 min (20-60k frames at 20-40 ms/frame) by leveraging the improved imaging kinetics afforded by DNA-PAINT-ERS^19^.

**Fig. 2:**
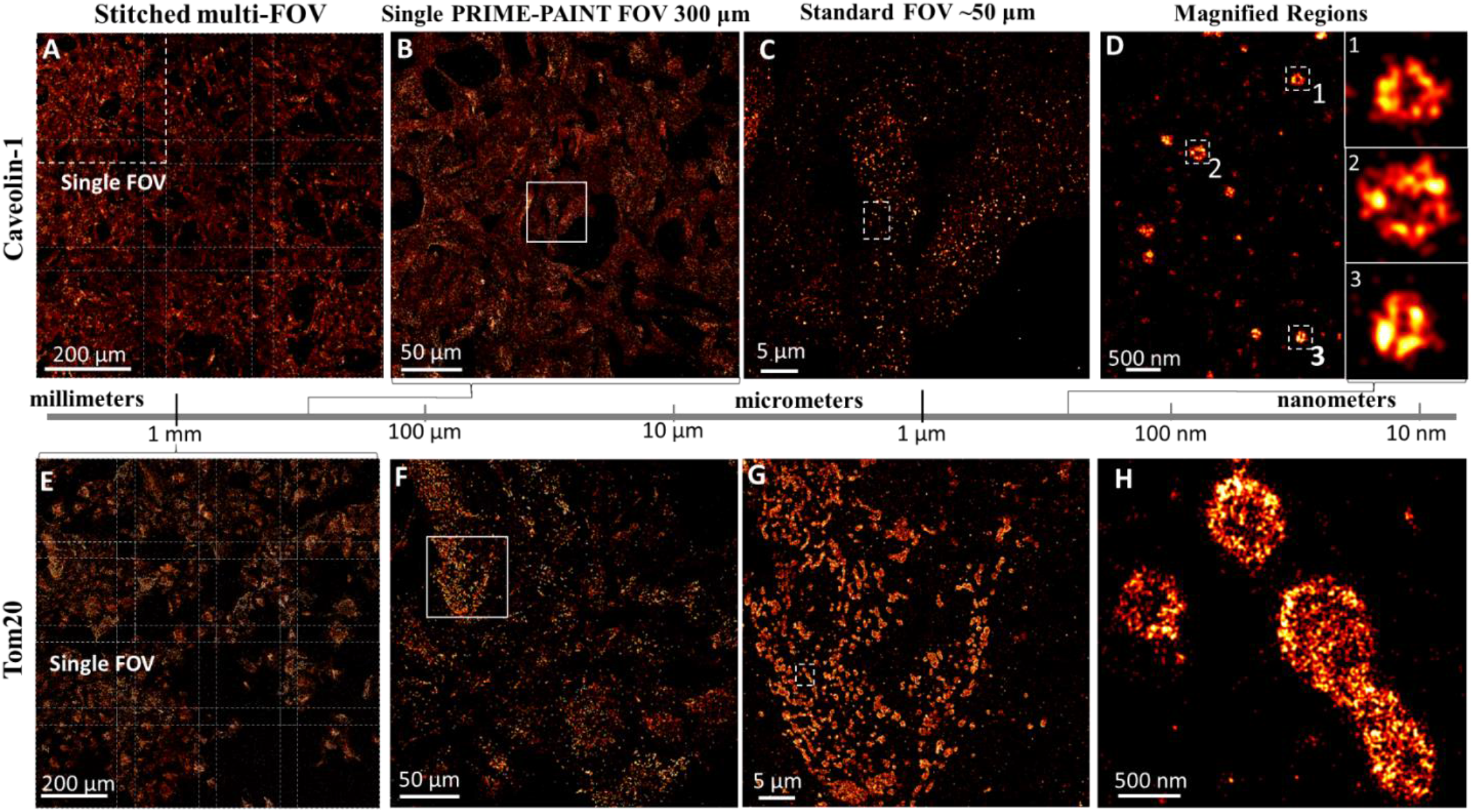
PRIME-PAINT imaging of caveolae and mitochondria over millimeter FOVs in U2OS cells. Representative single and stitched PRIME-PAINT images of membrane-adjacent caveolae (**A-D**) and cytosolic Tom20 (**E-H**). (**A**) Stitched 3x3 array of membrane-adjacent caveolae at 800 µm x 800 µm FOV with 40 µm overlaps. (**B**) Single PRIME-PAINT FOV at nearly 300 µm x 300 µm of Caveolin-1 imaged with strict TIR angle. (**C**) Magnified region from **B** highlighting the detail within a more standard smaller FOV of ~50 µm. (**D**) Magnified cell region from **C** and 3 insets showing high-quality super-resolution detail of caveolae vesicles. (**E**) Stitched 4x4 array of cytosolic Tom20 over 1 square millimeter. (**F**) Single PRIME-PAINT FOV at nearly 300 µm x 300 µm. (**G**) Magnified region from F highlighting mitochondria within a more standard smaller FOV of ~50 µm. (**H**) Magnified cell region from (**G**) with distinct mitochondria. Each single PRIME-PAINT image (**B**,**F**) was acquired in 15 minutes (30,000 frames at 30ms exposure) using 1 nM IS2-ATTO643 and 12.5% EC for both Caveolin-1 and Tom20. Stitched 3x3 FOV for caveolae was acquired in 135 minutes while 4x4 FOV for Tom20 was acquired in 240 minutes. Scale bars are 200 µm in (**A**,**E**), 50 µm in (**B**,**F**), 5 µm in (**C**,**G**), and 500 nm in (**D.H**). Insets in (**D**) are 300 nm wide.

The use of DNA-PAINT(-ERS) allowed robust stitching of multiple FOVs to address even larger sample areas. In **Fig. 2A** we demonstrate a 3x3 stitching (with a ~40 µm overlap) with a combined FOV of 0.8 × 0.8 mm^2^ in 1-2 hours when imaging caveolae, and **Fig. 2E**, a 4x4 stitching with a combined FOV of 1 mm^2^ in 2-3 hours and when imaging mitochondria (10-15 min per single FOV in each case). Leveraging the larger ~0.5 mm x 0.5 mm FOV enables imaging across ~1 mm^2^ as 2x2 stitches in under 40 minutes (**Supplementary Fig. S3**). Imaging of the same 1 mm^2^ FOVs would have taken 1-3 days if using DNA-PAINT with a standard 50 µm x 50 µm FOV or 6-8 hours even with ASTER^10^. Thus, PRIME-PAINT with DNA-PAINT-ERS^19^ substantially increases imaging throughput compared with existing strategies.

We estimated the effective resolution of PRIME-PAINT by measuring the width of microtubules. For these estimates, we assumed the width of microtubules to be around 40 nm (considering the inherent width and the size of the antibodies and the attached DS oligo). Microtubules exhibited Full Width Half Maximum (FWHM) of 47 ± 3 nm (objective-type DNA-PAINT-ERS) and 51 ± 4 nm (PRIME-PAINT at 0.3 mm x 0.3 mm FOV; see **Supplementary Fig. S4**). These correspond to an effective resolution of ~25 nm for the objective-type DNA-PAINT, consistent with our previous results^19^, and slightly lower (~30 nm) for PRIME-PAINT at 0.3 mm x 0.3 mm FOV using the Prime 95B camera. Imaging with the largest ~0.5 mm x 0.5 mm FOVs yield a lower lateral resolution of 40-45 nm (FWHM ~60 nm; **Supplementary Fig. S4**). The decrease in resolution is primarily due to the much-decreased laser power density, and a more powerful (2-3W; currently ~1W) laser should bring the resolution to the 20-30 nm range. In this work, we primarily used the 0.3 mm x 0.3 mm FOV to ensure optimal spatial resolution (~30 nm).

The integrated microfluidics (**Fig. 1C**) facilitated multiplexed PRIME-PAINT imaging by allowing manual or programmed buffer exchange. For example, we labeled microtubules, mitochondria, and vimentin in cultured COS7 cells and imaged the three targets sequentially using exchange-PAINT (**Fig. 3A & B**). A quick washing step between the imaging cycles by flowing in a blank imaging buffer (e.g. PBS with 15% EC) effectively eliminated any residual localizations within seconds (**Supplementary Fig. S5**). In the co-registered multitarget images (~0.3 mm x 0.3 mm FOV), all three targets were well resolved, revealing their spatial relationships such as occasional associations between mitochondria and the two cytoskeletal filaments (**Fig. 3C, D**). We similarly obtained a two-target PRIME-PAINT image of microtubules and vimentin over the larger, 0.5 mm x 0.5 mm FOV (**Fig. 3E-G**).

**Fig. 3:**
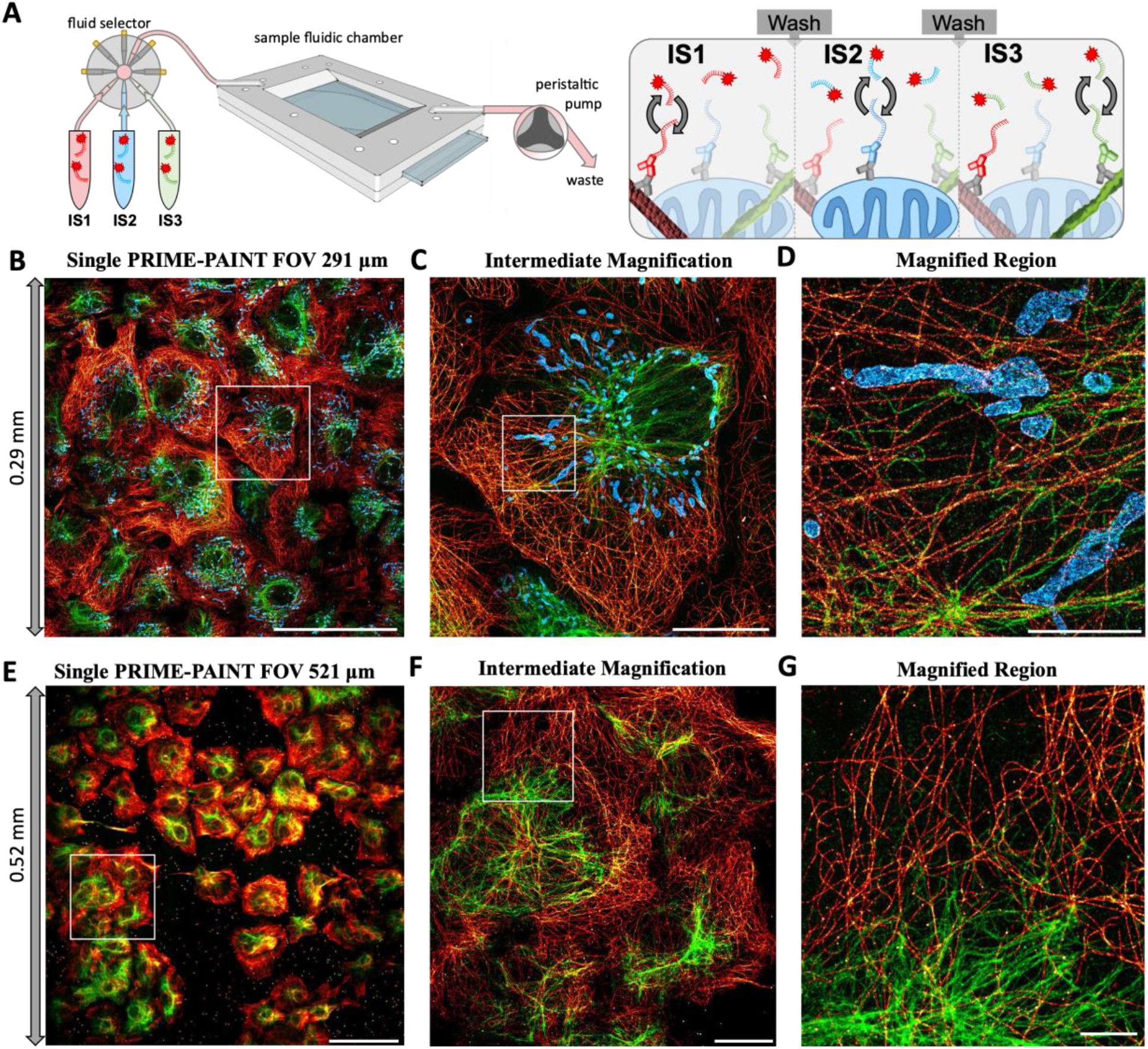
Multiplexed PRIME-PAINT imaging of cells via microfluidic exchange-PAINT. (**A**) Schematic of multiplexed imaging with complimentary imaging strand (IS) targeting docking strand (DS) to microtubules (red), mitochondria (blue), and vimentin (green). (**B**) Single PRIME-PAINT 3-target image of Cos7 cells at 300 µm x 300 µm. (**C**) Magnified region from **B** highlighting image quality over a more standard ~50 µm FOV. (**D**) Magnified region from **C** showing distinct cytoskeletal structures and mitochondria. (**E**) Expanded PRIME-PAINT 2-target FOV at 521 µm x 521 µm. (**F**) Intermediate magnification from **E** at 100 µm FOV. (**G**) Magnified region from **F** highlighting dense perinuclear vimentin and extended microtubule networks. The representative 3-target image (**B-D**) was acquired in 45 minutes total (30,000 frames at 30ms exposure for each target) using 1 nM IS1-ATTO643 and 12.5 % EC, 1 nM IS2-ATTO643 and 12.5 % EC, and 500 pM IS3-ATTO643 and 13.75 % EC for microtubules, mitochondria, and vimentin respectively. Expanded PRIME-PAINT FOV 2-target image (**E-G**) was acquired in 30 minutes total (30,000 frames at 30ms exposure for each target) using 1 nM IS1-ATTO643 and 12.5 % EC, and 500 pM IS3-ATTO643 and 13.75 % EC for microtubules and vimentin respectively. Scale Bars are 100 µm in (**B**,**E**), 20 µm in (**C**,**F**), and 5 µm in (**D**,**G**).

### Imaging and machine learning analysis of nanostructures in large cell populations

The high imaging throughput and resolution of PRIME-PAINT offers unique opportunities to analyze protein localizations and biological nanostructures across large cell populations. As a proof of concept, we analyzed caveolae in U2OS cells expressing KRas^G12D^, an oncogenic mutant of KRas. In initial observations, we found that cells expressing high KRas^G12D^ appeared to have fewer and/or smaller caveolae (**Figs. 4A-B**); we thus sought to further investigate this by leveraging the ability of PRIME-PAINT to quantitate the size and abundance of the caveolae in hundreds to thousands of cells at once.

**Fig. 4:**
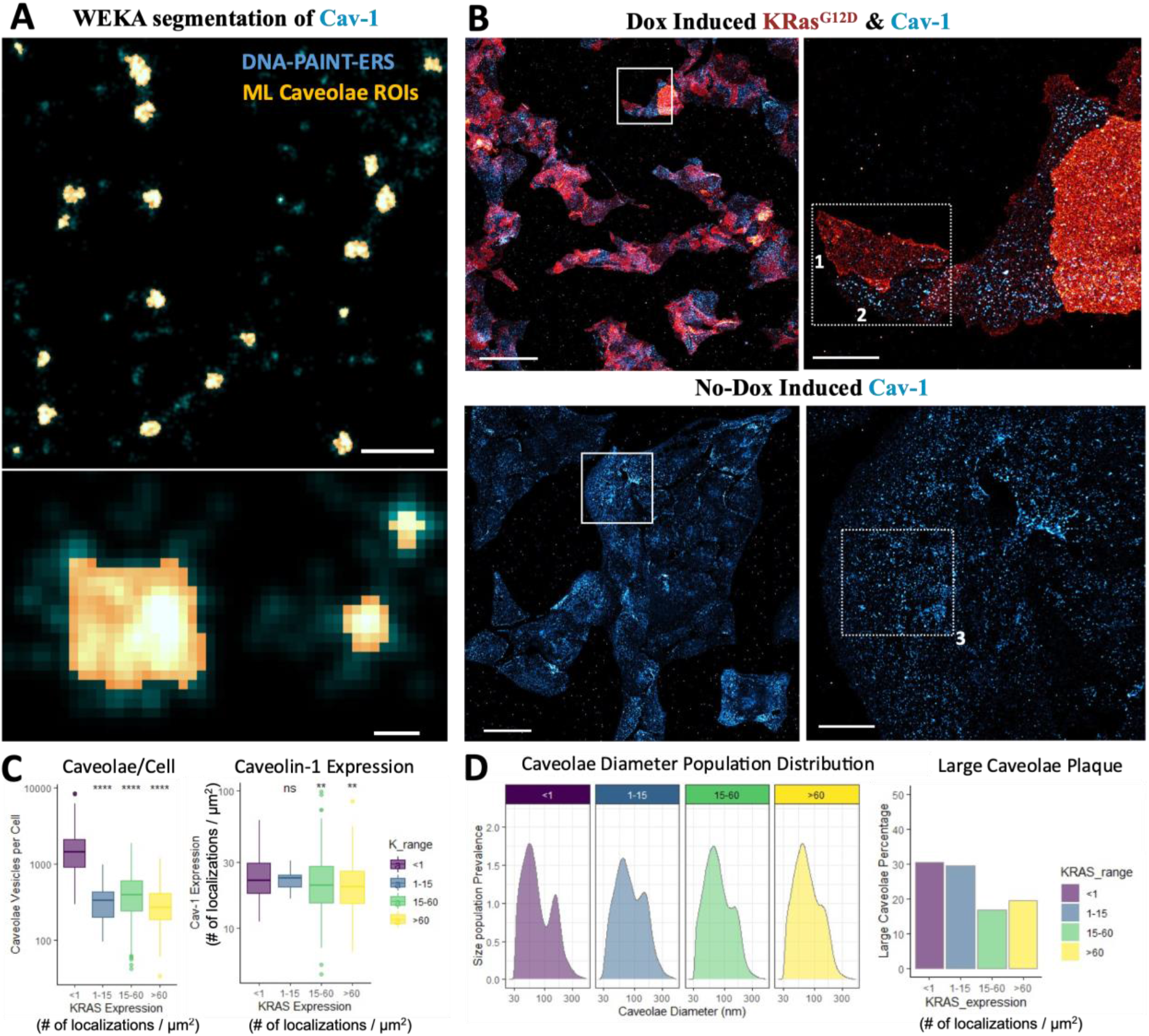
PRIME-PAINT image analysis of KRas G12D and caveolae in U2OS cells. (**A**) Representative overlays showing both caveolae vesicles as imaged via PRIME-PAINT in blue with corresponding caveolae outlines identified via machine learning segmentation in yellow (top). Inset showing both larger caveolae plaques and punctate caveolae vesicles, at ~150 nm and ~60 nm wide, respectively (bottom). (**B**) Representative PRIME-PAINT images of both dox-induced and non-dox induced KRas G12D and Caveolin-1. KRas G12D labeled via SNAP-tag DS-1 conjugate shown in red, with Caveolin-1 DS2 shown in blue. Magnified regions show representative cells numbered 1,2, and 3 with higher, lower, and no KRas G12D expressions respectively. Manually annotated cell boundaries were subsequently input into custom Fiji macro for both KRas G12D expression and WEKA classification of caveolae vesicles per cell. (**C**) Boxplot of the abundance of quantified caveolae vesicles versus ranges of mutant KRas expression (in units of # of localizations per µm^2^) for non-dox (<1), 1-15, 15-60, and >60 mutant KRas/um^2^. Boxplot of total Caveolin-1 expression versus ranges of mutant KRas expression per cell, with both expression values shown in # of localizations per µm^2^. (**D**) Population analysis of caveolae diameters for indicated KRas expression ranges, notably showing two main peaks at ~60 nm and ~150 nm from non-dox induction. Y-axis is the relative population of caveolae at differing diameters on the X-axis (log10 scaled). Higher KRas G12D expression anti-correlates with the larger caveolae size population. Analysis was performed on 10 images of dox-induced and two images of non-dox induced cells acquired with 30,000 frames at 30 ms exposure for each target, using 1 nM IS1-ATTO643 and 12.5 % EC, and 500 pM IS2-ATTO643 and 13.75 % EC for KRas G12D-SNAP-DS1 and Caveolin-1-DS2 respectively. Scale bars in (A) are 50 µm for full FOV images (left for each cell condition), 10 µm for matching magnified views (right), and 5 µm for all example cells (bottom panels). Plots in (**B**,**C**) were generated using ggplot package in R. Significance levels (in panel **B**) were calculated using wilcox test from average values within each cell relative to the non-dox induced cells group.

The U2OS cells were engineered to express SNAP-KRas^G12D^ upon doxycycline (dox)-induction^28^, where SNAP-KRas^G12D^ was labeled using DNA oligo-conjugated SNAP substrate (see *Methods*) and caveolae with antibodies as described earlier (**Fig. 2A-D**). We obtained and analyzed two-color images of mutant KRas and Caveolin-1 across 12 FOVs (~0.3 mm x 0.3 mm), 10 of which from Dox-induced (KRas^G12D^-positive) cells and 2 from uninduced cells (KRas^G12D^-negative). Next, we utilized WEKA^29^, an open-source platform for machine-learning with a built-in plugin for Fiji^30^, for automated identification and analysis of nanostructures from PRIME-PAINT images. As reconstructed PRIME-PAINT images were massive (~30,000 x 30,000 pixels at 10 nm/pixel rendering from a single ~0.3 mm x 0.3 mm FOV), we developed a Fiji macro to sub-divide, classify, and quantitate the nanostructures and recombine the quantitation results (**Supplementary Fig. S6**, also see *Methods*).

Using this strategy, we analyzed caveolae in PRIME-PAINT images of U2OS cells labeled for caveolin-1 (**Supplementary Fig. S7**). WEKA model performance was validated with a DICE similarity coefficient of 84% ± 8%. For each cell, we quantitated the cell area, signal intensity (i.e., total numbers of KRas and Caveolin-1 localizations), and nanostructure size (i.e., the diameter of individual caveolae) (**Supplementary Fig. S7 and Fig. 4A-B**). The level of expression for mutant KRas was calculated as the net number of KRas localizations (after combining localizations that span multiple raw frames) per µm^2^ image area. Collectively, 925 cells containing a total of ~630,000 caveolae were segmented and analyzed.

The analysis revealed an interesting relationship between caveolae size and KRas mutant expression. The number of caveolae per cell was significantly lower in KRas^G12D^ positive compared with KRas^G12D^-negative cells, confirming our initial observation. However, the total amount of caveolin-1 protein appears to not be affected by KRas^G12D^ expression, at least within the short-term (48-72 hours) Dox-induction used in this study (**Fig. 4C**). Importantly, at all KRas expression levels examined (0-60+ localizations/µm^2^), the histograms of caveolae diameter showed two distinct peaks at ~54 nm and ~154 nm (**Fig. 4D**), which we tentatively attributed to fully formed caveolae (~60 nm diameter) and precursor ‘plaques’ (~150 nm diameter) as previously identified in EM images^31^. As KRas^G12D^ expression increased, the peak diameters remained largely unchanged (**Supplementary Fig. S8**) but the fraction of large caveolae noticeably decreased (**Fig. 4D**). This observation may support that active KRas promotes caveolae maturation, echoing previous reports^32,33^, although the effect may be cell-line and context specific.

While a full validation of this result and its mechanistic interpretation warrant a more extended study, this example demonstrates the feasibility in utilizing PRIME-PAINT in combination with machine-learning based image segmentation in probing alterations in nanoscopic structures across large cell populations, which would have been difficult if not for both the nanoscale resolution and the larger FOVs.

### Microfluidics-enhanced DNA-PAINT nanoscopy of clinical FFPE tissue samples

Although FFPE tissue nanoscopy via STORM has been demonstrated for years^17^, reports demonstrating the same with DNA-PAINT have been lacking. The main motivations for imaging FFPE tissue sections with DNA-PAINT would be the high spatial resolution, convenient multiplexing, and reliable stitching. In DNA-PAINT, photobleaching is limited to the fluorophore attached to the imaging strand oligo and does not affect the target-attached docking strand. As such, imaging of the current FOV has little impact on the neighboring sample areas (**Supplementary Fig. S9A-C**), permitting robust stitching across as many FOVs as necessary to address even larger sample areas.

To our surprise, initial attempts of DNA-PAINT imaging on FFPE tissue sections yielded a much lower localization density compared to that on cells (**Supplementary Fig. S9D-E**) and, in consequence, poorly reconstructed images, despite efficient and on-target labeling as confirmed by examining the signals from Cy3 attached to the DS oligos. Considering the quality difference with previous tissue STORM results^17^ prepared using nearly identical antigen retrieval and similar staining methods, we suspected that the lack of DNA-PAINT signals was due to inefficient DS-IS hybridization. Accidentally, we observed that the localization kinetics was drastically improved when the perfusing pump was left on during image acquisition (**Supplementary Fig. S10A**). Follow-up investigations showed that a weak flow at 1 µL/min could already increase the observed localizations by ~10-fold and resulted in significantly better image quality (**Supplementary Fig. S10B-C**). No additional benefits were evident when the flow rate was increased to 5µL/min. In contrast to imaging the tissue sections, microfluidic flow during imaging only situationally helped DNA-PAINT imaging of cells grown on glass, perhaps because the localization kinetics was already near optimal on cultured cells (**Supplementary Fig. S11**).

Several other, empirical modifications to sample preparation and imaging steps helped further improve PRIME-PAINT imaging of FFPE tissue sections. First, we found that the optimal EC% for FFPE tissue sections is between 5-8% (for cells typically 10-15% EC) (**Supplementary Fig. S12**). This might be related to refractive index matching between the EC-containing buffer and the sample^34^. Second, addition of Signal Enhancer (SE), a charge-blocking commercial product previously used in our DNA-PAINT(-ERS) sample preparations^19^, followed by RNase A treatment during blocking prior to antibody incubations yields best PRIME-PAINT images on FFPE sections (**Supplementary Fig. S13**).

### Multiplexed nanoscopy of pancreatic tumor sections with PRIME-PAINT

We chose to use FFPE sections of Pancreatic Ductal Adenocarcinoma (PDAC) to exemplify multiplexed tissue nanoscopy with PRIME-PAINT. Dual labeling for Tom20 and pan-cytokeratin could inform structural changes in mitochondrial organization, which are theorized to occur during PDAC development^35–38^. To capture such structural changes with fine details, nanoscopic imaging of whole pre-PDAC and PDAC lesions – often hundreds of µms across – is necessary.

A histological overview photographed with a 20x objective revealed a moderately differentiated PDAC within desmoplastic stroma (**Fig. 5A & B**). Using signals from Cy3-DS conjugated to the secondary antibodies as a reference, immunofluorescence confirmed strong pan-cytokeratin staining along the duct and faint mitochondria both in and around the duct (**Fig. 5C**). By acquiring six adjacent FOVs with PRIME-PAINT at ~0.3 mm x 0.3 mm each with a 40 µm overlap, we imaged the entire duct and proximal stroma covering an area over 800 µm x 500 µm (**Fig. 5D**). At an acquisition speed of 30 minutes per FOV per target, the entire imaging took ~3 hours for each target and ~6 hours for a dual-target image.

**Fig. 5:**
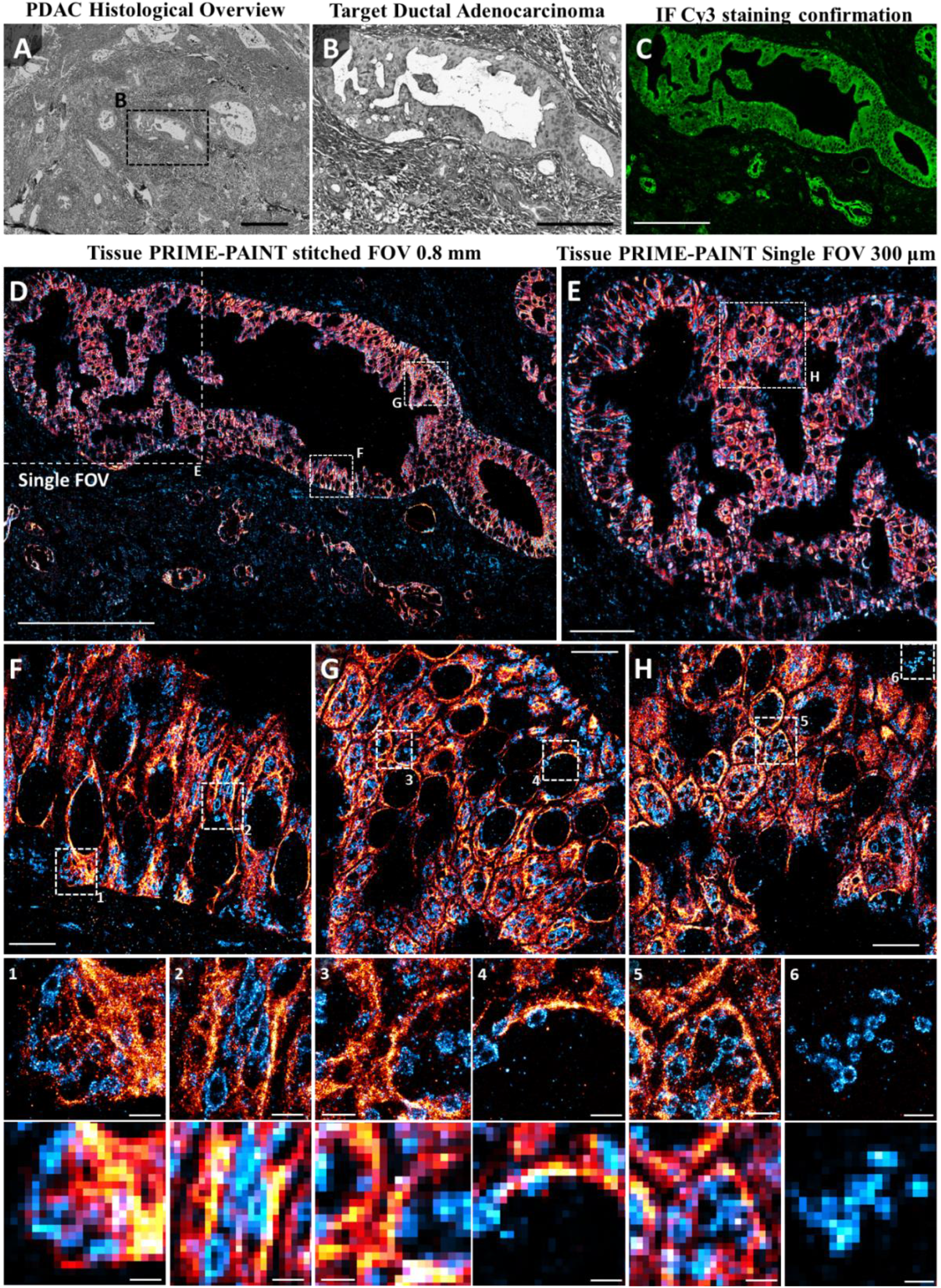
Multiplexed PRIME-PAINT of pancreatic cancer tissue. (**A**) Histological overview of moderately differentiated PDAC within desmoplastic stroma acquired at 20x magnification. (**B**) Targeted ductal adenocarcinoma for imaging with PRIME-PAINT. (**C**) Immunofluorescent confirmation of Cy3 signal from secondary antibodies conjugated to docking strand oligos showing strong pan-cytokeratin staining along the tumor and diffuse mitochondrial labeling within the tumor and adjacent stroma. (**D**) Stitched tissue PRIME-PAINT image of an entire, 800 µm long ductal adenocarcinoma with both prognostic pan-cytokeratin in red and mitochondrial Tom20 in blue. (**E**) Single tissue PRIME-PAINT image obtained under mild flow (‘microfluidics-enhanced’). (**F-H**) Select magnified regions from (**D**,**E**) highlighting the high quality imaging of cellular features seen within different regions of the tumor. Numbered insets shown with matching diffraction limited views (300 nm pixel size). Tissue PRIME-PAINT 2-target images (**E-G**) were acquired in 6 hours total (30,000 frames at 60 ms exposure for each target) using PRIME-PAINT at 1 µL/min flow and 500 pM IS1-ATTO643 and 7 % EC, and 500 pM IS2-ATTO643 and 7 % EC for pan-cytokeratin and Tom20 respectively. Scale bars are 500 µm in (**A)**, 200 µm in (**B-D**), 50 µm in (**E**), 10 µm in (**F-H**), and 2 µm for both PRIME-PAINT and diffraction-limited bottom numbered insets.

Close-up views demonstrate the uniform image quality over the entire FOV (**Fig. 5E**). In smaller ROIs, sub-diffractive details of mitochondria and pan-keratin can be seen in unparalleled detail versus an otherwise diffraction limited blur (**Fig. 5F-H**). Not only can distinct mitochondria of varying sizes and shapes be seen, but dense pan-cytokeratin also tapers to fine, filamentous networks often enclosing the cell periphery. Finally, to further exemplify the full range of spatial information obtainable by PRIME-PAINT on FFPE tissue sections, we serially magnified a second ductal adenocarcinoma from the millimeter-scale overview down to distinct mitochondrial membrane structures (**Supplementary Fig. S14**). Based on this multiscale view of entire tissue regions, we should be able to quantitatively compare the structure and spatial organization of mitochondria and the intermediate cytoskeleton in different types of cells within the same tissue as well as in tissues from different origins (such as lesions at different stages of PDAC).

## Discussion

We have demonstrated PRIME-PAINT, a new platform for multiplexed bioimaging across roughly five orders of length scales (~10 nm to mm scales). PRIME-PAINT is based on a simple optical design and capitalizes improvements in imaging kinetics, sample preparation, and microfluidics for fast and high-quality biological nanoscopy of broad sample types.

Although there are already several solutions available for large-field biological nanoscopy, PRIME-PAINT has some unique advantages. PRIME-PAINT affords among the largest single imaging FOVs (~0.5 mm x 0.5 mm) of all fluorescence nanoscopies, comparable to that of chip-based nanoscopy. It also offers among the best lateral resolution (40-45 nm) at this FOV and ~30 nm at the smaller, ~0.3 mm x 0.3 mm FOV, comparable to previous nanoscopy modalities with through-objective imaging schemes.

As such, PRIME-PAINT allows imaging of millimeter scale sample regions in as little as 40 minutes per target. Lastly, PRIME-PAINT is built on DNA-PAINT and thus naturally excels at multiplexed imaging, especially with the integrated microfluidic sample chamber. These traits make PRIME-APINT ideally suited for studying complex biological systems with multiscale and multi-component features.

The prism illumination scheme has been widely used in single-molecule imaging studies. The imaging setup reported in this work, which is slightly modified from the original prism-type TIR, comprises only off-the-shelf components and can be readily adopted without needing sophisticated scanning optics^10^ or microfabricated imaging substrates^12^. The integrated microfluidic imaging chamber can also be constructed using only commercial parts, although a custom, machined chamber holder is encouraged for improved mechanically stability and reliability. As such, PRIME-PAINT can be constructed and maintained for routine operations in most biology laboratory settings.

We expect that PRIME-PAINT can be readily extended in several aspects. The spatial resolution of PRIME-PAINT was primarily limited by the excitation power density and could be further improved with simple solutions such as using a more powerful laser or adopting an ‘ASTER’-like scanning strategy^10^. PRIME-PAINT is also amenable to automation through integration of programmable fluidics, FOV stitching and buffer exchange^39^, and on-line data processing^40,41^. Additionally, leveraging recent progress in multiplexing strategies^42–44^ and 3D calibration over extended depth and large FOVs^45,46^, 3D PRIME-PAINT imaging of many more targets over several mm^2^ sample areas should be readily feasible.

By significantly extending the imaging throughput of fluorescence nanoscopy, PRIME-PAINT and the other recent, large-FOV modalities^9–12^ bring exciting new opportunities. Similar to cryo-EM single particle analysis, which revolutionized structural biology, recent work has used nanoscopy data to infer dynamic protein complexes^47,48^. Large scale nanoscopic image datasets will synergize with machine-learning-based image analysis tools such as WEKA^29^ and LocMoFit^49^ to significantly accelerate biomedical discovery^50,51^. Additionally, the ability to image FFPE sections at nanometer resolution across mm^2^ FOVs (**Fig. 5 and Fig. S14**) is an important step toward clinical applications, for which PRIME-PAINT and the likes represent a leap forward compared with prior attempts using electron microscopy (EM)^52–54^ or fluorescence nanoscopy with a small FOV^16,17^.

## Materials and Methods

### Materials

Methods utilized for generating labeling reagents followed established protocols for DNA-PAINT-ERS ^19^. In brief, all starting DNA oligonucleotides were obtained from Integrated DNA Technologies. Docking strands included a 5′ amino modifier C6, for further conjugation with DBCO-PEG4-NHS (Click Chemistry Tools, A134-2) via succinimidyl ester chemistry, and a 3′ Cy3™ fluorophore which helps confirm proper antibody labeling. Imaging strands with a 3’amino group were reacted using succinimidyl ester chemistry with NHS-ATTO 643 (ATTO-TEC, AD 643-31). All reactions were performed at at room temperature in ultrapure water adjusted to pH~8.5 using 1 M sodium bicarbonate (Fisher Scientific, M-14636) for 3 h. Conjugated DNA oligos were purified via ethanol precipitation, and resuspended in Invitrogen™ UltraPure™ DNase/RNase-Free Distilled water (Thermo Fisher Scientific, 10977023).

Secondary antibodies used were: AffiniPure Donkey anti-Rabbit IgG (H+L) (Jackson Immuno Research 711–005–152), AffiniPure Donkey anti-Mouse IgG (H+L) (Jackson Immuno Research 715–005– 150) and AffiniPure Donkey anti-Chicken IgG (H+L) (Jackson Immuno Research 703-005-155); these were conjugated with azido-PEG4-NHS via succinimidyl ester chemistry. Antibody-PEG4-azide conjugates were purified through a 50 kDa Millipore Sigma™ Amicon™ Ultra Centrifugal Filter Unit (Fisher Scientific, UFC505096). Next, purified antibody-PEG4-azide were reacted with excess DBCO-DS (molar ratio 1: 5) via copper-free click chemistry, overnight at room temperature using a rocker. Antibody-DS products were isolated through a 100 kDa Millipore Sigma™ Amicon™ Ultra Centrifugal Filter Unit (Fisher Scientific, UFC510096). Protein concentrations and the degrees of labeling were found using the peak signals at 260, 280 nm, 550 nm (for Cy3™) in a Nanodrop UV-Vis spectrophotometer (ThermoFisher Scientific, 2000c). In general, oligo-conjugated secondary antibodies generated contained 4-5 conjugated DS oligos.

The SNAP-tag substrate BG-PEG4-Azide was synthesized by the Medicinal Chemistry Core at Oregon Health and Science University. Briefly, it was synthesized in two steps via an amine-reactive key intermediate prepared from commercially available BG-NH2 as the starting material, followed by an NHS-ester crosslinking reaction. The final crude product was purified by preparative HPLC. The structure and purity of BG-PEG4-Azide were further confirmed by analytical HPLC analysis and high-resolution mass spectrometry prior to the click oligonucleotide labeling. BG-PEG4-Azide was reacted with excess DBCO-DS (molar ratio 1:10) via copper free click chemistry, on a rocker at room temperature overnight. The resulting BG-DS was purified via ethanol precipitation, and suspended in UltraPure™ DNase/RNase-Free Distilled water. The concentration was determined by the Nanodrop UV-Vis spectrophotometer, similarly as mentioned above.

Primary antibodies used were: Mouse-beta tubulin monoclonal antibody (Thermo Fisher Scientific, 32–2600), Rabbit-anti-Tom20 polyclonal antibody (Abcam, ab78547), Rabbit-anti-caveolin-1 antibody (Abcam, ab2910), Chicken-anti-vimentin antibody (Sigma Aldrich, AB5733), and Mouse panCytokeratin (Abcam, ab7753).

For all remaining experimental steps and sample processing and labeling (Fixation, permeabilization, and immunostaining, etc) materials used included: paraformaldehyde (Sigma-Aldrich, P6148), Triton X-100 (Sigma-Aldrich, X100), 25% glutaraldehyde (Sigma-Aldrich, G6257), bovine serum albumin (Fisher Scientific, BP1600), sodium hydroxide (Fisher Scientific, S318-100), sodium borohydride (Sigma-Aldrich, 452882), Invitrogen™ Salmon Sperm DNA (Thermo Fisher Scientific, AM9680), sodium azide (Fisher Scientific, AC190381000), Gibco™ Dulbeccos PBS with calcium and magnesium (PBS+) (Thermo Fisher Scientific, 14–040–182), and 50 nm gold particles (BBI Solutions, EM.GC50/4). Fixation was performed using a buffer made from: 2x PHEM buffer, generated with 0.06 M PIPES (Sigma-Aldrich, P6757), 0.025 M HEPES (Fisher Scientific, BP310-500), 0.01 M EGTA (Thermo Fisher Scientific, O2783-100), and 0.008 M MgSO4 (Acros, 4138–5000) in distilled water, 10 M potassium hydroxide (Sigma-Aldrich, 221473) was used to finally adjust the pH to 7. As with DNA-PAINT-ERS, a % volume combination of EC (Sigma-Aldrich, 676802) with buffer C (PBS plus 500 mM Sodium Chloride) was used as indicated for both imaging and even washing steps.

### Flow-chamber preparation and cell culture

Flow chamber substrates were made using: 25x75mm fire-polished microscope slides (Schott, Nexterion® Slide Glass B 1025087). Fire-polished microscope slides were each drilled twice using a 1/16th inch bit diamond coated drill bit (Lasco Diamond #F6) for later use as microfluidic inlet and outlet ports, at the coordinates of (4 mm, 16 mm)- and (21 mm, 59 mm) on the 25x75mm coverslide. After drilling, slides were rinsed with DI water (3x), and sonicated in 100 % EtOH for 10 minutes. Following 3x DI water rinses, slides were etched in 1M NaOH for 20 minutes. After three rinses with DI water, cleaned slides were left in 100% EtOH prior to cell culture/tissue slide preparation.

### Cell and Tissue samples

Cell lines used in this study included U2OS (ATCC®, HTB-96) and COS7 (ATCC® CRL-1651™). U2OS and Cos7 cells were passaged every 3–4 and 2–3 days respectively, and cultured in Gibco DMEM (Thermo Fisher Scientific, 11995073) mixed with 10% fetal bovine serum (Thermo Fisher Scientific, 26–140–079). Passaging was performed using Trypsin-EDTA (0.25%) (Thermo Fisher Scientific 25200056), with cells kept to below 15 passages. For SRM imaging experiments, cells were grown on custom drilled coverslides within a sterile oval silicon cutout until 50-60% confluency prior to fixation.

All patient FFPE tissue samples (including HER2+ breast cancer and PDAC samples) were collected through either the OHSU Biorepository or the Brenden Colson Center following IRB approved protocols including patient consent for research applications. Standard human pancreas samples obtained by BCC were FFPE sections from otherwise healthy cadavers and were used for most tissue imaging optimizations. FFPE tissue samples were cut using a ultramicrotome in 2 πm thick sections (RM2125 RTS, Leica Biosystems, Germany).

Pre-drilled and cleaned fire-polished coverslides were prepared for tissue mounting using Poly-Ethylene-Imine (PEI) (Sigma--Aldrich, 904759-100G) coating. In brief, cleaned coverslides were treated for 20 minutes with 0.1% PEI solution in ultrapure H2O. After coating, excess PEI was rinsed 3x with ultrapure H2O for 5 minutes each. After aspirating excess H2O, coated slides were left to dry flat at 42°C for 2+ hrs, or until completely dried. Fluidic chamber profile outlines were drawn onto the non-PEI-coated backside using a ultrafine sharpie to assist in tissue mounting positioning. All tissue samples were sectioned at 2um ±0.5um thickness, floated on a 42°C waterbath immediately prior to mounting onto PEI-coated coverslides. Mounted tissues sections were dried vertically for 1 hour at 60°C before storage.

Mounted tissue sections were stored vertically in this manner for up to 1 month prior to antigen-retrieval, labeling and imaging.

### Immunostaining

For immunostaining of caveolin and SNAP, cells were fixed for 20 min with 3.7% paraformaldehyde (PFA) in 1x PHEM buffer, after a quick PBS wash. Following two PBS washes, cells were quenched with fresh 0.1% sodium borohydride in PBS for 7 min, and followed 3 washes with PBS (5min each). Cells were permeabilized with 0.3% saponin in PBS for 20 min. For immunostaining of microtubules, Tom20 and vimentin, cells were fixed for 20 min with 3.7% PFA and 0.1% glutaraldehyde (GA) in 1x PHEM, followed by 3x PBS washes, quenching with sodium borohydride and permeabilization in 0.2% Triton X-100 in PBS. Blocking in PBS with 3% bovine serum albumin, 5% salmon sperm DNA (Thermo Fisher Scientific, AM9680) for 45 min was done on a rocker, followed by incubation with the primary antibody for Tom20 (1:250), caveolin (1:200), tubulin (1:100) or vimentin (1:250) antibodies in PBS buffer containing 3% BSA. The incubation took place overnight on a rocker at 4°C in a humidity chamber. Next, cells were washed three times (5 min each) with PBS before incubation with respective secondary antibody-DS described above at a final concentration of ~8 µg mL^−1^ in PBS buffer containing 1% BSA and 5% salmon sperm DNA; the secondary antibody incubation also took place on a rocker at room temperature for 90 min. For DS secondary antibody incubation and subsequent steps, the sample was kept in the dark to avoid bleaching of conjugated fluorophores. Cells were washed three times with PBS (5 min each). All cell samples were post-fixed for 10 min with 3.7% PFA and 0.1% GA in 1x PHEM. Before imaging, cells were incubated with 15% 50 nm gold particles in PBS+ for 1 min, followed by a quick PBS wash.

For immunostaining of FFPE tissue samples, FFPE sections were deparaffinized using xylene (2x, 10 min), 100% EtOH (2x, 10 min) 95% EtOH in DI water (5 min), 70% EtOH in DI water (5 min), 50% EtOH in DI water (5 min) and left in PBS. Tissues underwent antigen retrieval in a decloaking chamber (Bio SB, BSB-7087) first in Tris buffer (300mM Tris, 0.05% Tween 20, PH 8). Tissues were transferred into a pre-heated citrate buffer (300mM NaCitrate Monohydrate, 0.05% Tween 20, PH 6), also heated during decloaking, and allowed to cool to room temperature. After two PBS washes, tissues were further permeabilized with 0.4% Triton X-100 in PBS for 45 minutes. After removing excess permeabilization solution and three PBS washes, a hydrophobic barrier (company product) was applied around the mounted tissue following the fluidic outline of the chamber design. Tissues were either further treated with RNAseA/ Image-iT® FX signal enhancer (ThermoFisher Scientific, I36933) or directly blocked prior to antibody labelling. In brief, RNAse A and signal enhancer treatment both occurred prior to antibody labeling as optional optimizations. After permeabilization, tissue samples could be treated with RNAse A (ThermoFisher Scientific, EN0531) at 50x dilution in PBS on a rocker for overnight at room temperature. After a rinse with PBS, sections were quenched with fresh 0.1% sodium borohydride in PBS for 7 min, and followed by three washes with PBS (5min / each). Tissue was further incubated with signal enhancer at room temperature for 30 minutes, followed by three washes with PBS wash (5 min/each).

After PBS washes, tissues were blocked with 3% BSA and 0.3% saponin in PBS for 1 hour. Next, tissues were incubated with primary antibodies: Tom20 (1:200 dilution), and panCytokeratin (1:150 dilution) in PBS containing 3% BSA and 5% salmon sperm DNA. The incubation took place on a rocker overnight at 4°C in a humidity chamber. Following three PBS washes (5 min each), tissues were incubated with respective DS-conjugated secondary antibodies at a final concentration of ~8 µg mL−1 in PBS buffer containing 1% BSA and 5% salmon sperm DNA. The incubation also took place on a rocker at room temperature for 2 hours, followed by three PBS washes (5 min each). After which, all tissue samples were post-fixed by 3.7% PFA and 0.1% GA in 1x PHEM at room temperature for 30 min. Before flow chamber assembly, tissues were incubated with 15% 50 nm gold particles in PBS+ for 1 min, followed by a quick PBS wash.

### Flow Chamber assembly

Flow chamber exterior was made using a CNC-cut aluminum holder which fits gently outside the sample sandwich, providing compression to meet matching 1/16 fractional width O-rings (McMaster-Carr, 2418T11) and Tygon tubing (Cole-Parmer, EW-06419-01) at the pre-drilled points on the coverslide. This external holder and microfluidic tubing can be easily reused for each sample (see .CAD file, **Supplementary data**). Since the sample coverslide still provides the backbone of the fully assembled sample and fluidics for imaging, both the microscope objective and prism can be used at nearly identical positions between samples.

To complete flow chamber interior assembly around immunolabeled cells/tissues mounted to pre-drilled coverslides, 24 mm x 40 mm high-precision #1.5 coverslips (Thorlabs, CG15KH), and 2-inch wide Behmis® Parafilm (Thermo Fisher, 12-374-16) were first combined. After sonicating coverslips in 100 % EtOH for 10 minutes, coverslips were quickly air dried using a gentle stream of compressed air. Parafilm was cut into 1 inch wide sections and stretched length-wise until just prior to tearing. Stretched Parafilm was placed onto coverslips such that no folds or air-pockets were present. Excess Parafilm was removed from the coverslip border using a razorblade, and fluid profile stencil was used to cut the flow chamber interior into the parafilm. After peeling the inner part of the Parafilm, the coverslip and fluidic profile were ready for final assembly. After removing excess PBS from cells/tissues, coverslips with fluidic profiles were slowly placed on top of the coverslide with sample such that no excess solution contacted the Parafilm border. A 110 W glue gun tip (with glue removed) was used to gently press the coverslip along the Parafilm border to melt the Parafilm and seal the fluidic chamber.

Fluidic components used were MX Series II™ 10 Position/11 Port Selector Valve (IDEX Health & Science, MXX778-605) and RP-TX Peristaltic Pump (Takasago Electric, RP-TXP5S-P04A-DC3VS). Pump control was achieved through Arduino Uno (Amazon, X7375-10G) and Adafruit Motor/Stepper/Servo Shield for Arduino v2 Kit (Adafruit, 8541582581). Both the port selector and peristaltic pump were controlled via custom C code.

## Microscopy

Through-objective TIRF superresolution data in this work were taken on a custom single-molecule imaging system described previously ^19^. Briefly, two lasers emitting at 561 nm (Opto Engine LLC, 150 mW), and 647 nm (Coherent OBIS 647, 140 mW) were combined and introduced into the back of a Nikon Ti-U microscope equipped with a 60× TIRF objective (Nikon, Oil immersion, NA 1.49). An f = 400 mm lens was placed at the back port of the microscope to focus the collimated laser light to the back aperture of the objective to achieve objective TIR illumination. The excitation light can be continuously tuned between epi-fluorescence and strict TIR angle modes by shifting the incident laser horizontally with a translational stage before entering the back port of the microscope. Additionally, a weak cylindrical lens compensates for longitudinal beam widening through the prism, to maintain power density over a round FOV for efficient single-molecule excitation. A custom focus stabilizing system based on detection of the reflected excitation laser was used to stabilize the focus during data acquisition. A multi-edge polychroic mirror (Semrock, Di01-R405/488/561/635) was used to reflect the lasers into the objective and clean up fluorescence signals from the sample. Emission filters used for the 561 nm (for imaging Cy3 on the DS), and 647 nm (for imaging ATTO643 conjugated ISs) were FF01-605/64 and FF01-708/75, respectively (all from Semrock). Fluorescence signals were collected through the objective by an electron-multiplied charge-coupled device (EM-CCD, Andor, iXon Ultra 897) using a typical EM gain setting at 200–300 in frame transfer mode. Unless otherwise indicated, the power density of the 647 nm laser (for DNA-PAINT imaging using ATTO643 conjugated IS) was typically around ~500 W/cm^2^.

Prism-type TIRF PRIME-PAINT data was collected using a custom single-molecule imaging system as outlined in Fig. 1A. Key system components used from the laser to the camera were: 1W 639 nm laser (Opto Engine, MRL-FN-639-1W), 2x beam expander (ThorLabs BE02M-A), cylindrical lens f=1000 mm, 5x beam expander (ThorLabs GBE05-A), 2-5x continuous beam expander (ThorLabs BE02-05-A), 4π lens cage (using an f = 100 mm and an f = 80 mm lens), 10 mm Square Aperture UV Fused Silica Prism (Pellin Broca, ADBU-10), 40x silicon oil objective (Nikon, Plan APO 40x/1.25 Sil λS WD 0.3, MRD73400), 647nm long pass filter (Semrock, LP02-647RU-50), 633 nm Stopline® notch filter (Semrock NFD01-633-25x36), and finally either a side-mounted Prime-95B sCMOS (Teledyne Photometrics, Prime 95B 25MM) or Kinetix sCMOS (Teledyne Photometrics). All detection path components housed within a Nikon Eclipse Ti2-E microscope body. Perfect Focus Unit (Nikon PFS, TI2-N-NDA-P) was integrated into the microscope body and provided stable autofocusing even during mild flow integration during PRIME-PAINT imaging.

### Data acquisition and image processing

Superresolution images were acquired using the open source micromanager software suite (https://micro-manager.org/)^55^ and saved as OMERO TIF files. Image analyses for extracting single-molecule localization and subsequent localization filtering, sorting, and rendering was performed using in-house Matlab scripts^56^. Briefly, raw localizations were first filtered based on localization fitting parameters such as signal to noise ratio, widths of point spread functions in the x and y dimensions, aspect ratio, etc. Next, the localizations were sorted, during which events that appeared within a defined number of frames (typically 2–3) and distance (typically 80 nm) were then combined into a single event with averaged coordinates. The sorted localizations were then used for final image rendering, and the rendered images were saved as TIF files for further analysis and annotations in Fiji. Multi-color images were co-registered using an average of observed gold fiducial positions within each FOV, while multi-FOV stitched images were aligned using average position of observed overlapping gold fiducials along shared image boundaries.

### Custom Machine-learning integrated Fiji Macro

Our analysis Macro was built using the Fiji macro language in .ijm format. Machine learning segmentation/classification was performed using Trainable WEKA Segmentation plugin within Fiji ^29^. Training was performed using input images with manually drawn caveolae particle boundaries trained against background diffuse cytoplasmic caveolin-1. Subsequent testing of trained model performance against manually annotated caveolae on three additional datasets resulted in a DICE coefficient of 84±8%.

Macro workflow is visually shown in Supplementary Fig. S7. This begins by opening a multitarget PRIME-PAINT rendered image, typically 29,100 x 29,100 pixels, and using Fiji ROI manager to manually draw boundaries for each cell of interest to quantify. After initiation, the full PRIME-PAINT image is loaded into the WEKA plugin and the trained .model file for caveolae is loaded, with all subsequent steps happening automatically. For each picked cell ROI, the macro will cut the cell out of the full PRIME-PAINT image, masking signal outside the ROI, and automatically generate sub-images with an adjustable overlap to tile across each cell ROI analyzed. Each sub-image is temporarily saved, and WEKA classification is performed on each cell’s sub-image. After classifying each sub-image of a particular cell, dynamic offsets initially added during sub-image generation are removed and the full classified cell is combined using original coordinates. Distinct ROI for all detected vesicles are saved and used to measure original cell image attributes at the positions indicated from WEKA classification outputs. In practice, our custom Fiji macro would automatically segment caveolae from input cells at a rate of ~30 minutes per cell, although this performance will vary depending on the computer specifications used.

These per-cell caveolae measurements are saved as .csv outputs and can be post-processed using programs such as R.

### Population analysis and other plots

Outside of rendered images, all plots were generated in R using the ggplot package. Caveolae particles smaller than 20 nm were excluded from downstream analysis as their border could not be reliably determined and were not considered as a part of initial manual annotation for observed caveolae particles. Population analysis was performed using the nls() function in R, which was used to model a two gaussian population within the total observed populations of caveolae particle sizes according to diameter.

### Statistical analysis

Image analysis was performed using custom semi-automated machine learning macro installed in ImageJ (see **Supplementary Fig. S7-8**). DNA-PAINT images with manually annotated cell boundary .roi were input to extract caveolae particle positions as .roi lists. Mapping these per-caveolae particle .roi positions onto the original image, the ImageJ “Measure” function was used to extract and save .csv files per cell, with image attributes including Area, Diameter, Mean intensity, etc. All downstream analysis and visualization was performed in R. P-value calculations in Fig. 4C were performed using the Wilcoxon test in the stat_compare_means() function within the ggplot package in R. For Fig. 4D, the Y axis of size population prevalence calculated using the geom_density function in ggplot from log_10_-transformed X axis of all caveolae particle diameters.

## Acknowledgments

The authors thank Drs. Joe W Gray, Gordon Mills, Terry K Morgan, Young Hwan Chang, Jason Link, Sean Speese, Yu-Jui (Roger) Chiu, David Qian, Kai Tao, and many other colleagues at OHSU for their helpful discussions. M.J.R, F.C., J.K., D.H., T.Z., Y.Z., S.E., and X.N. are members of and supported by the Cancer Early Detection Advanced Research (CEDAR) Center of the OHSU Knight Cancer Institute. The work is supported in part by the OHSU Knight Cancer Institute, the Damon Runyon Cancer Research Foundation, the Cancer Systems Biology Consortium from the National Cancer Institute (CSBC, grant number U54 CA209988, PI: Joe W. Gray), and the National Institute of General Medical Sciences (grant number R01 GM132322, PI: X.N.).

## Supplementary materials

Please see attached Supplemental information (.pdf containing supplementary figures S1-14) and Supplementary software (.zip file containing CAD files for the custom parts).

## Author contributions

Conceptualization: XN, FC, MJR; Methodology: MJR, XN, JK, DH, TZ, MS, JS; Investigation: MJR, JK, DH, YZ, TZ; Visualization: MJR, JK, YZ; Supervision: XN, SE; Writing—original draft: MJR, XN; Writing—review & editing: MJR, XN, MS, TZ, YZ, SE

