## Supplementary material for "Multiplexed and millimeter-scale fluorescence nanoscopy of cells and tissue sections via prism-illumination and microfluidics-enhanced DNA-PAINT"

**This PDF file includes:**

Figs. S1 to S14

**Additional Supplementary data includes:**

Data S1: Fluidic holder design

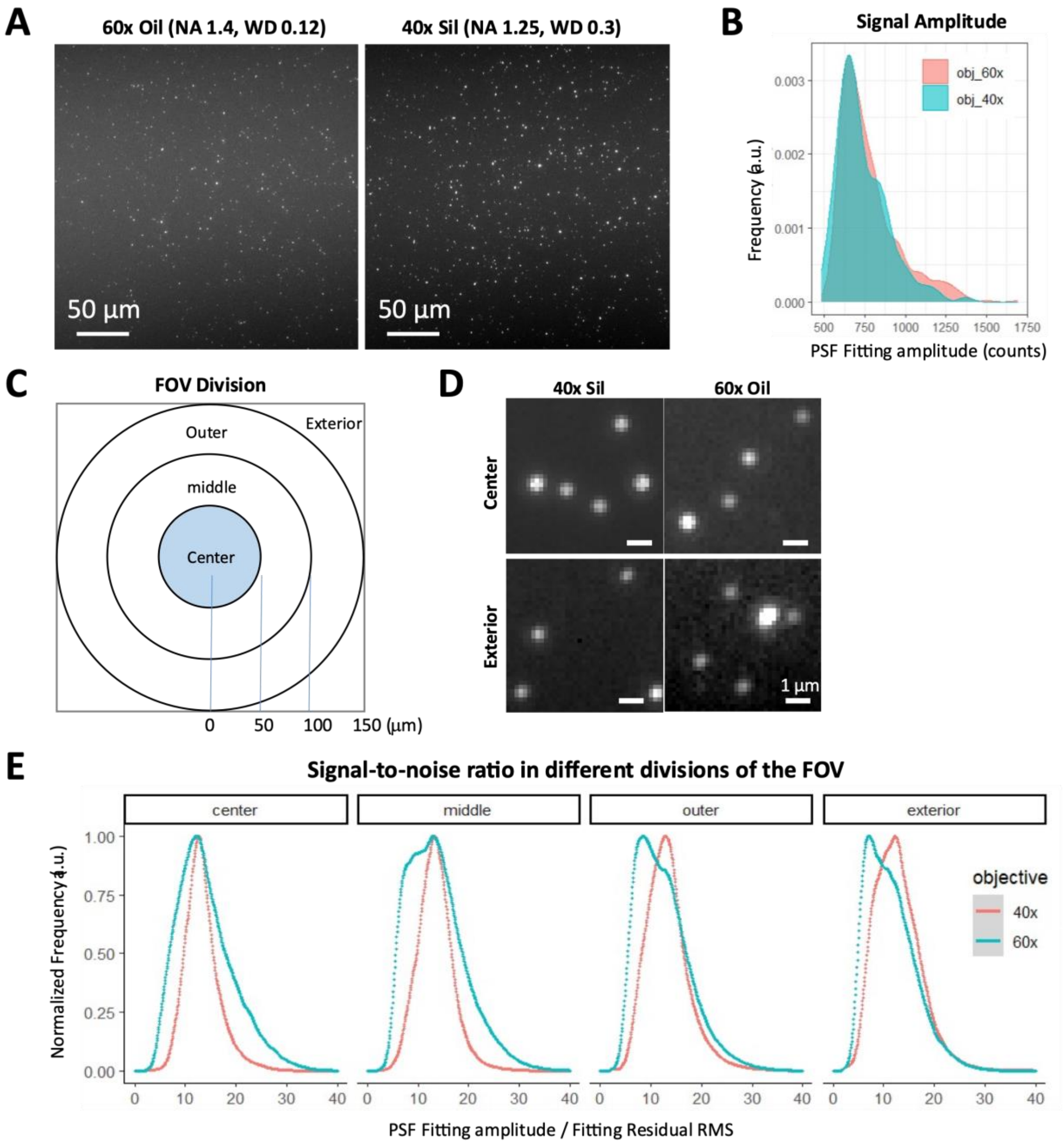


**Fig. S1.**

Comparing the performance for PRIME-PAINT between a 60x oil immersion (Nikon, WD 0.12, NA 1.4) and a 40x silicon oil (Nikon, WD 0.3, NA 1.3) objective. (A) Images of 50 nm gold nanoparticles deposited on the coverslide, sandwiched, and imaged similarly to a regular cell sample using the 60x (left) or 40x (right) objectives. Images shown using the same pixel intensity ranges; (B) Histograms of the PSF fitting amplitude of all nanoparticles imaged using the two objectives; (C) Dividing the FOV radially into four regions for comparing the single particle image quality as shown in (D) as raw images and (E) in terms of single-to-noise (defined as PSF fitting amplitude / fitting residual root mean square (RMS)) ratios.

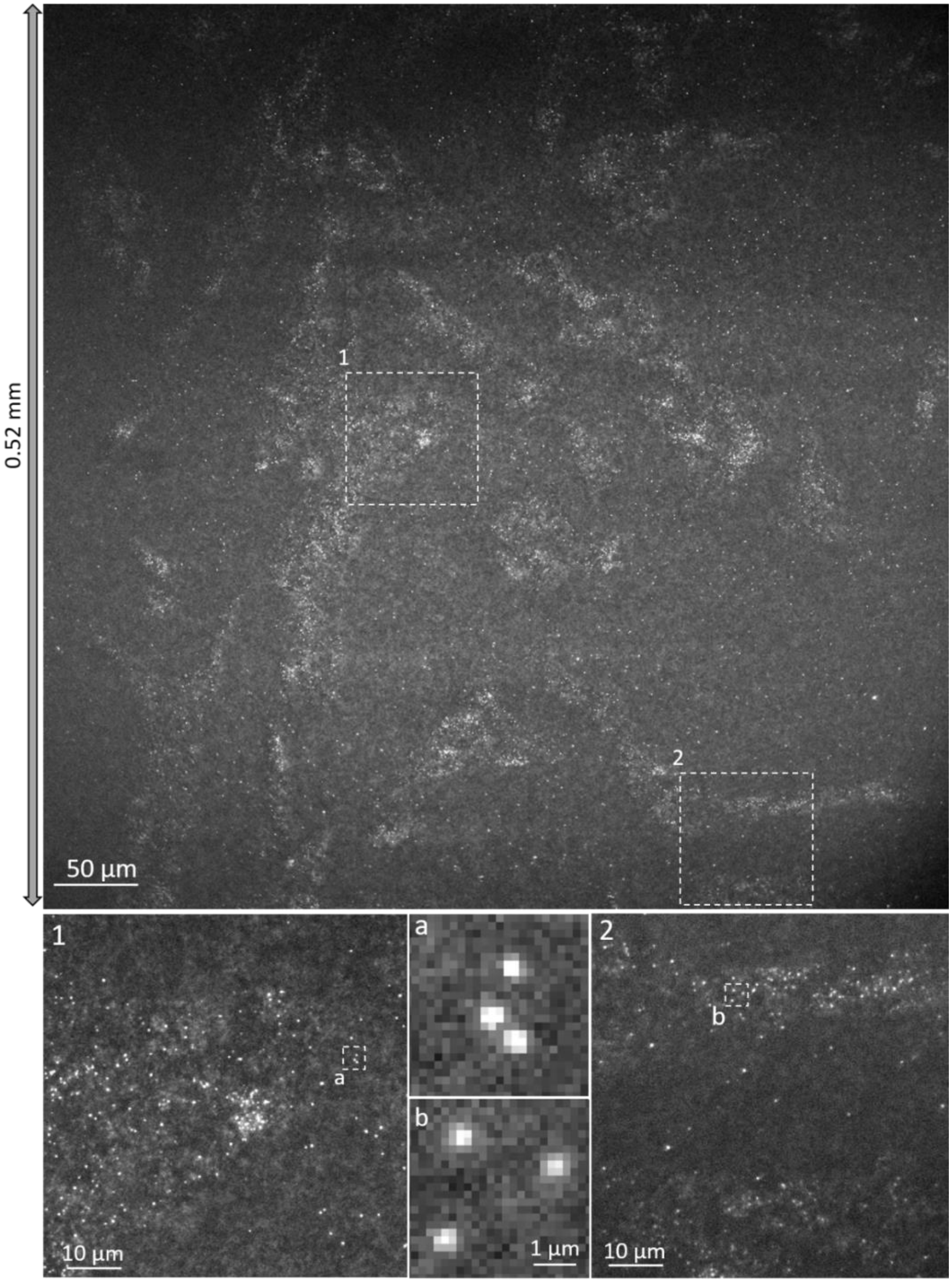


**Fig. S2.**

Representative expanded 521 µm x 521 µm PRIME-PAINT raw image taken using Kinetix camera at 40 ms exposure, highlighting single-molecule localization quality. Single-molecule events are clearly resolved both in the center on the FOV, and near the corners, exemplifying the potential to expand PRIME-PAINT FOVs to even larger areas.

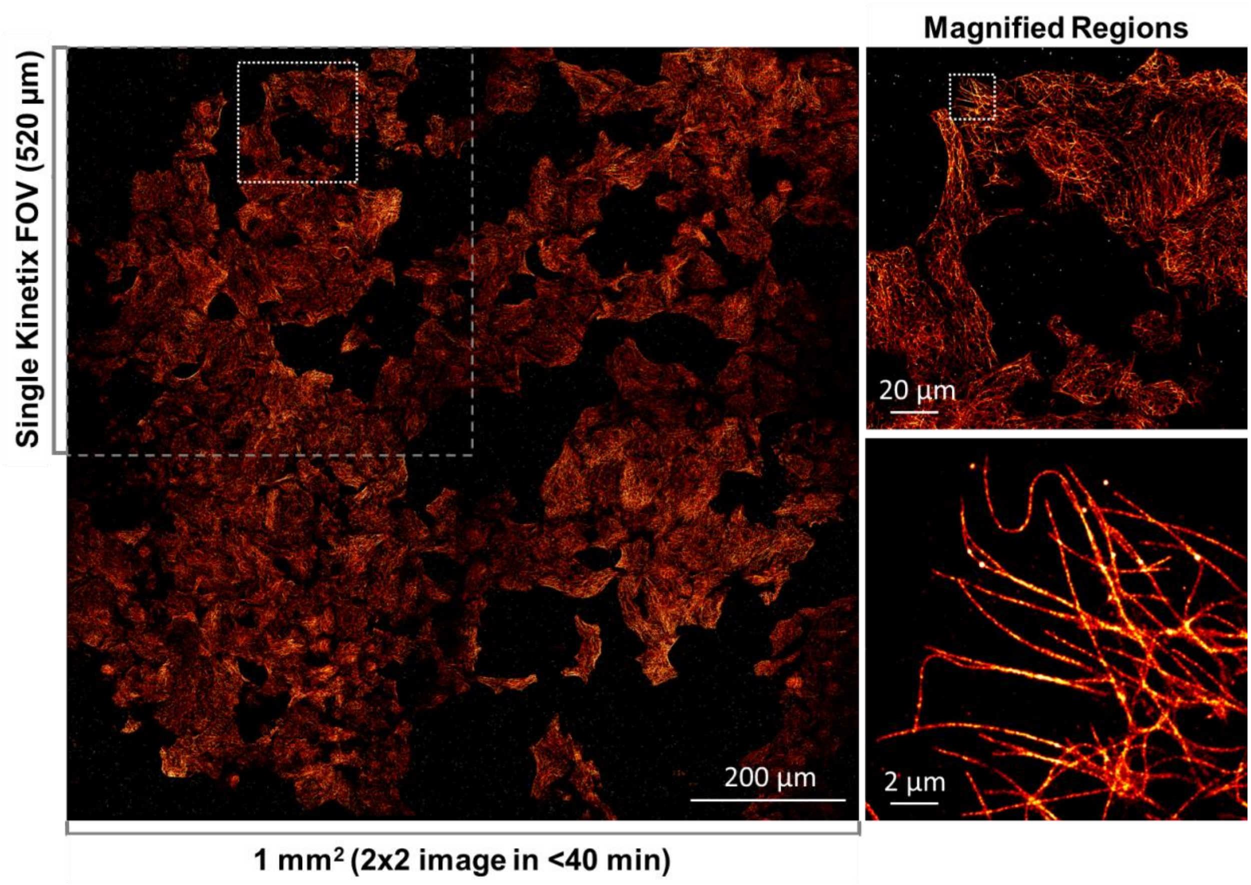


**Fig. S3.**

Representative stitched PRIME-PAINT 2x2 array of microtubules images at 1 mm x 1 mm FOV with 40 µm overlaps. Magnified regions show multiple adjacent cells as well as microtubules within one cell. Each single PRIME-PAINT image collected using the larger Kinetix sCMOS camera was acquired in 8 minutes (15,000 frames at 30ms exposure) using 1 nM IS1-ATTO643 and 12.5% EC. Stitched 2x2 full FOV was acquired in < 40 minutes. Scale bars are 200 µm (left panel), 20 µm (upper panel) and 2 µm (bottom panel).

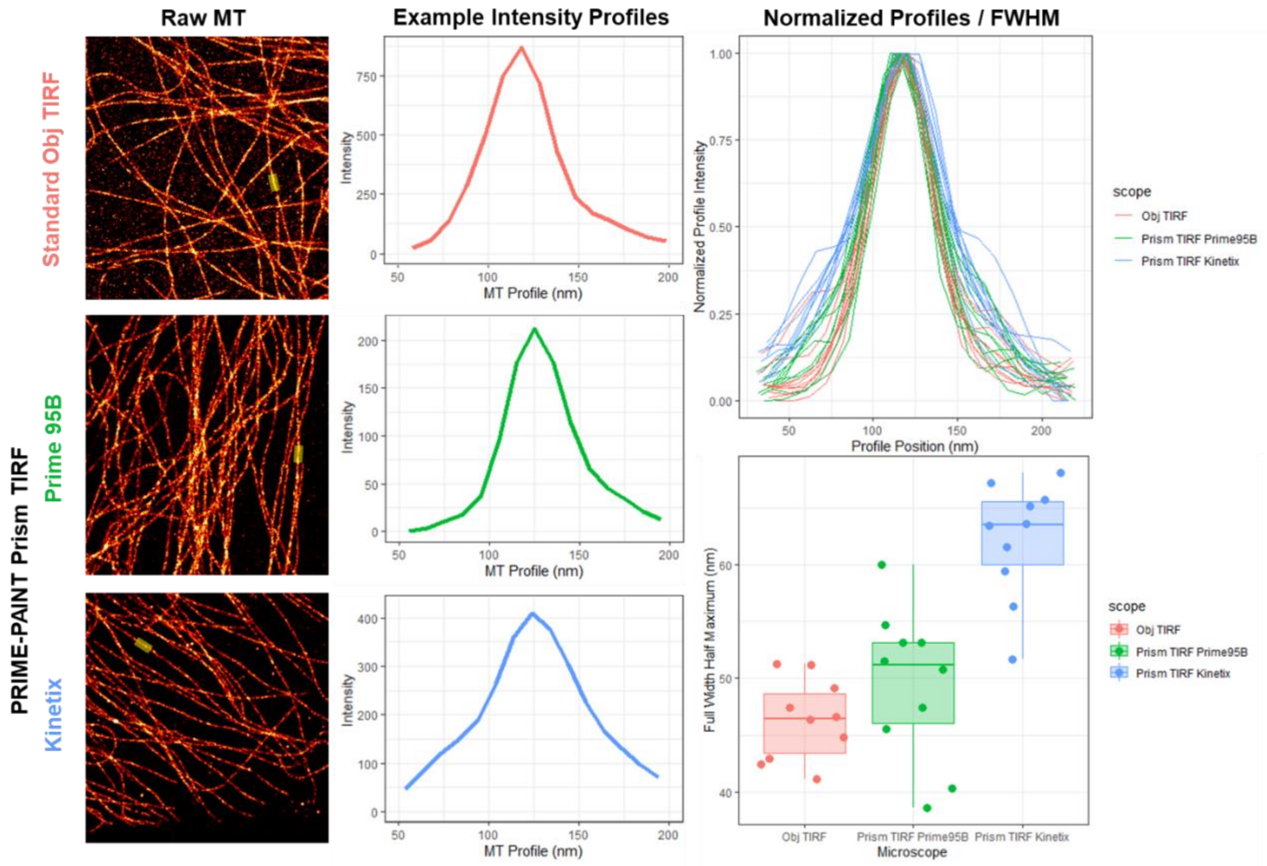


**Fig. S4.**

Comparable resolution of microtubules imaged via our standard objective-type TIRF microscope (Obj TIRF) and our PRIME-PAINT prism-type TIRF setup (0.3 mm Prime95B and 0.5 mm Kinetix camera FOVs). Representative reconstructed images of microtubules from both microscopes (left column), and example intensity profiles (middle column). Normalized intensity for 10 line profiles from both microscopes/cameras and calculated Full-Width Half-Maximum (FWHM) resolution (right column). All images were acquired with 30,000 frames at 30 ms exposure using 1 nM IS1-ATTO643 and 12.5 % Ethylene Carbonate (EC).

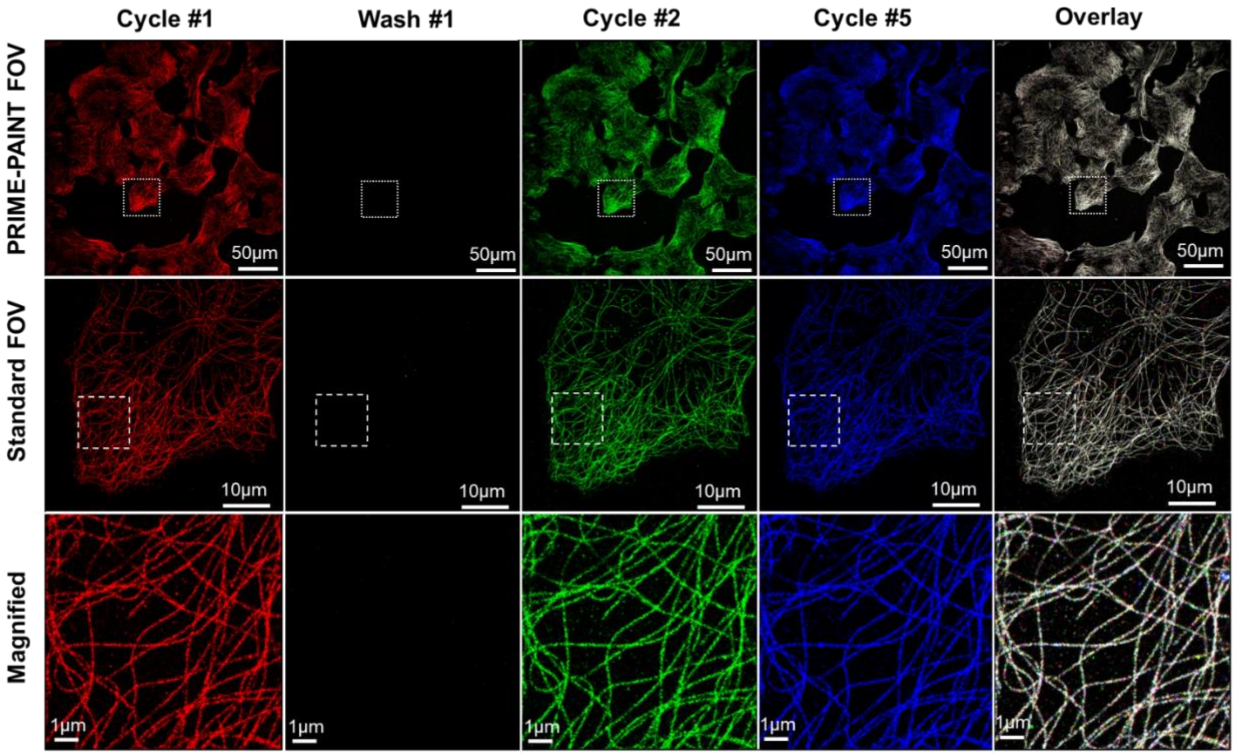


**Fig. S5.**

Reproducible microfluidic exchange and image quality of Cos7 cell microtubules using PRIME-PAINT. Representative views from 1st, 2nd, and 5th cycles with intermediate 15 % EC washes shown in red, green, and blue, respectively. Overlay of the three indicated cycle colors shows high similarity converging onto white microtubules. Full PRIME-PAINT FOVs (top row) with matching views of a single cell within a smaller, more standard FOV of 50 µm (middle row). Final magnified views highlight microtubule quality (bottom row). Scale bars are 50 µm (top row), 10 µm (middle row) and 1 µm (bottom row).

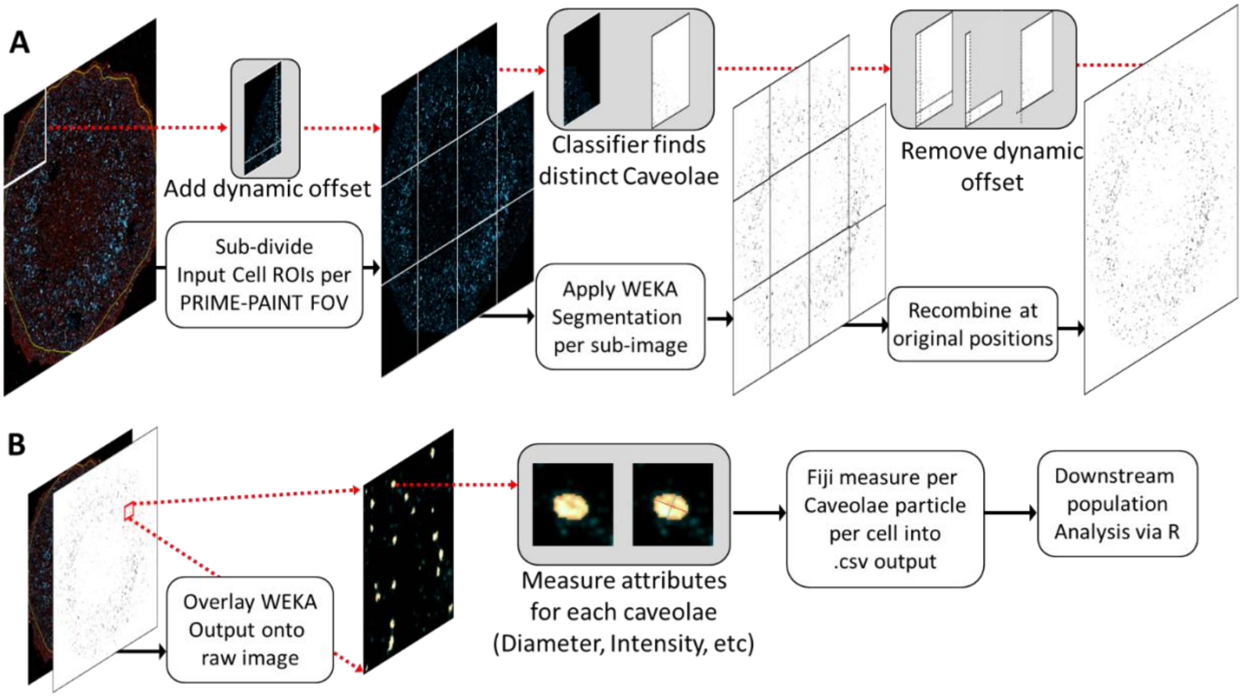


**Fig. S6.**

Schematic workflow of our custom Fiji macro integrating WEKA segmentation. (A) Representative steps for a single input cell ROI: 1) sub-divide total input cell ROI into smaller squares dynamically and add an offset (dynamic means: this will auto-adjust the number of squares and corresponding offsets within predefined upper bounds for square sizes in order to fill different size cell ROIs), 2) WEKA segmentation is applied serially to each sub-image using optimized classifier .model file for caveolae classification, 3) dynamic offsets are removed (to avoid edge effects) and full cell input with classified caveolae is recombined at original sub-image positions. (B) Caveolae feature extraction workflow: 1) recombined WEKA output per cell (from A) is overlaid back onto input cell PRIME-PAINT reconstructions, 2) using this mask, individual caveolae ROI are saved and used to measure image attributes such as diameter, mean intensity, etc, using Fiji *measure* function, 3) aggregate measurements per cell and per PRIME-PAINT FOV are saved into .csv files for downstream processing and visualization using R. Representative input cell shown was from the dox-induced KRAS$^{G12D}$ datasets as quantified in Figure 4.

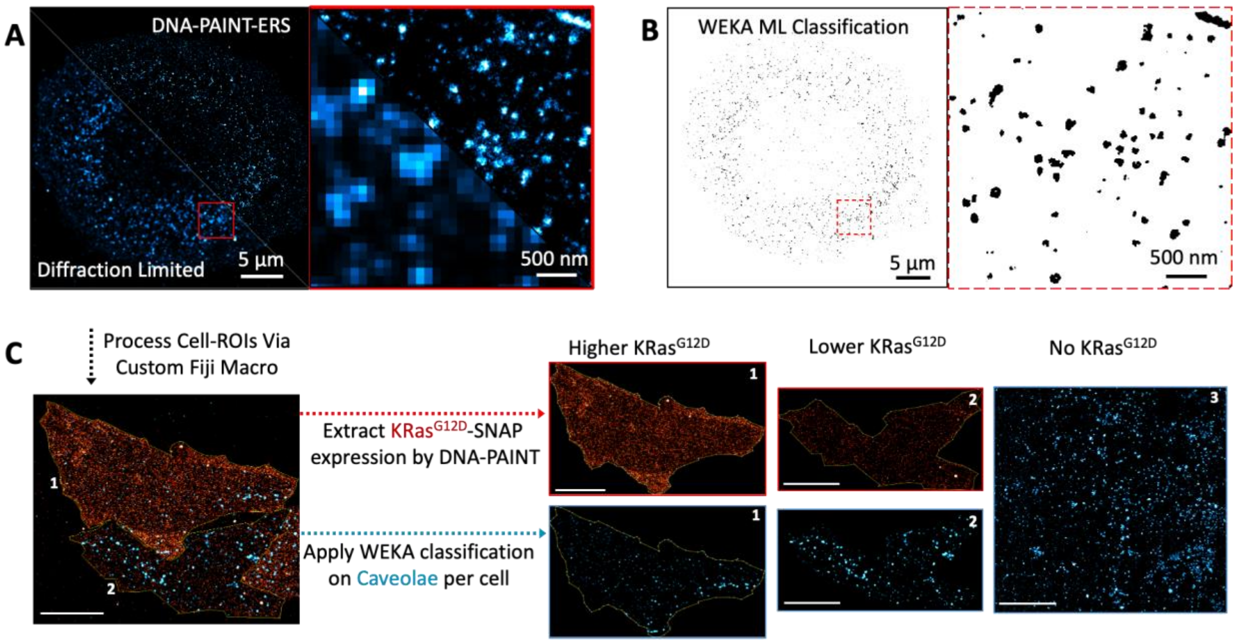


**Fig. S7.**

Schematic workflow of analyzing KRas expression levels and caveolae structure using the custom Fiji macro and trained WEKA segmentation. (A) PRIME-PAINT image of caveolae in U2OS cells, showing the whole cell (left) and zoom-in views (right, top right half) along with the corresponding diffraction-limited image (right, bottom left half); 2) Individual caveolae identified via WEKA segmentation, yielding Fiji ROIs that are then analyzed to extract the number, size and other properties; (C) Combining KRas expression level (per cell based on the net number of localizations) and WEKA segmentations of caveolae for investigating the impact of KRas G12D expression on caveolae abundance and structure.

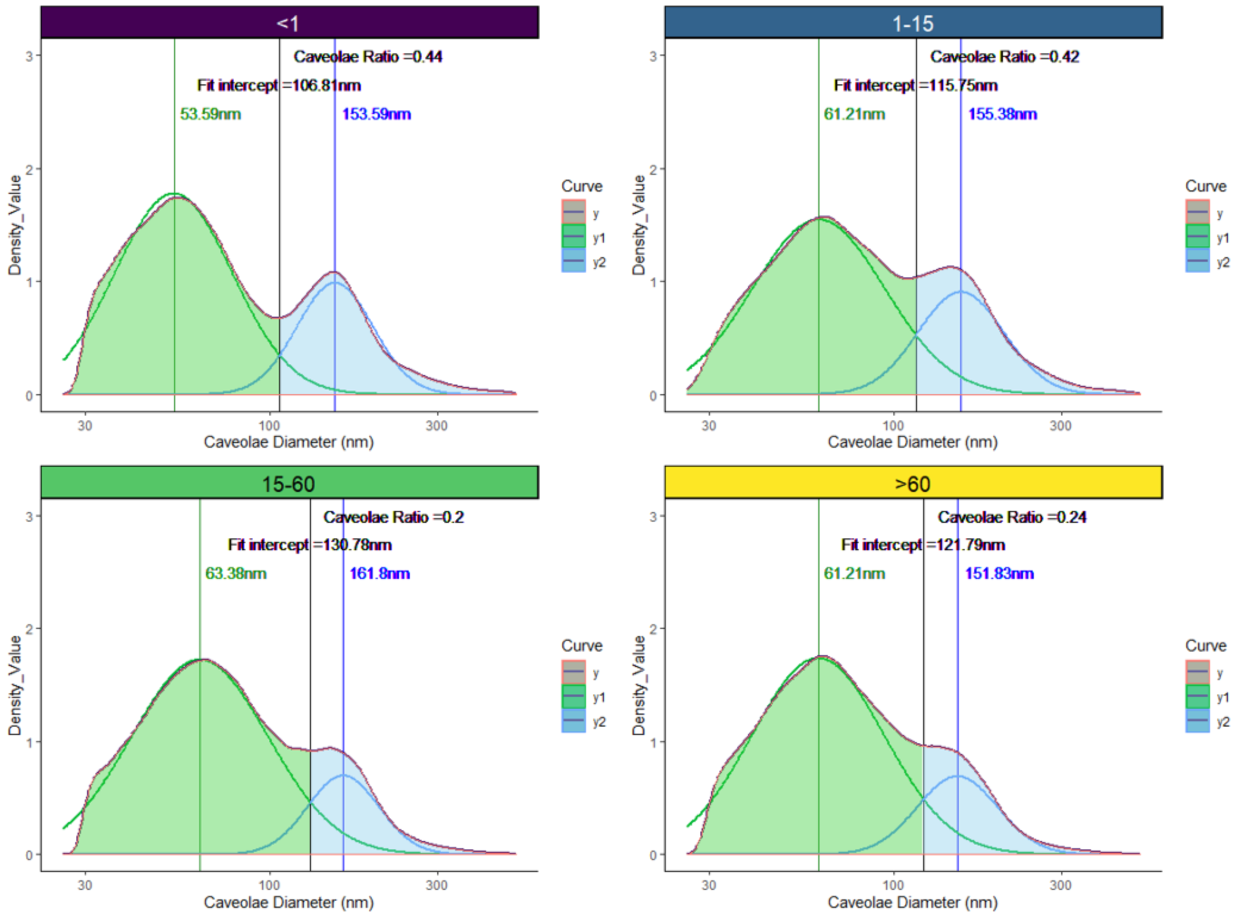


**Fig. S8.**

Population analysis of caveolae vesicles from dox-induced $KRAS^{G12D}$ experiment (Figure 5). Density plot of caveolae diameter (in nm) shown per indicated KRAS expression ranges: (<1, i.e. non-dox induction), (1-15), (15-60), and (>60) KRAS / $\mu m^2$. The R function *nls()* was used to fit two gaussian curves within the larger total population distribution resulting in a prominent smaller population size with peak ~60 nm (green), and larger population peak at ~155 nm (blue). The intercept of these two population curves was used to approximate the relative contribution of both size ranges within the total caveolae population observed for each expression range of $KRAS^{G12D}$. The indicated caveolae ratio shows the relative abundance of the larger caveolae population area (blue) divided by the smaller population area (green).

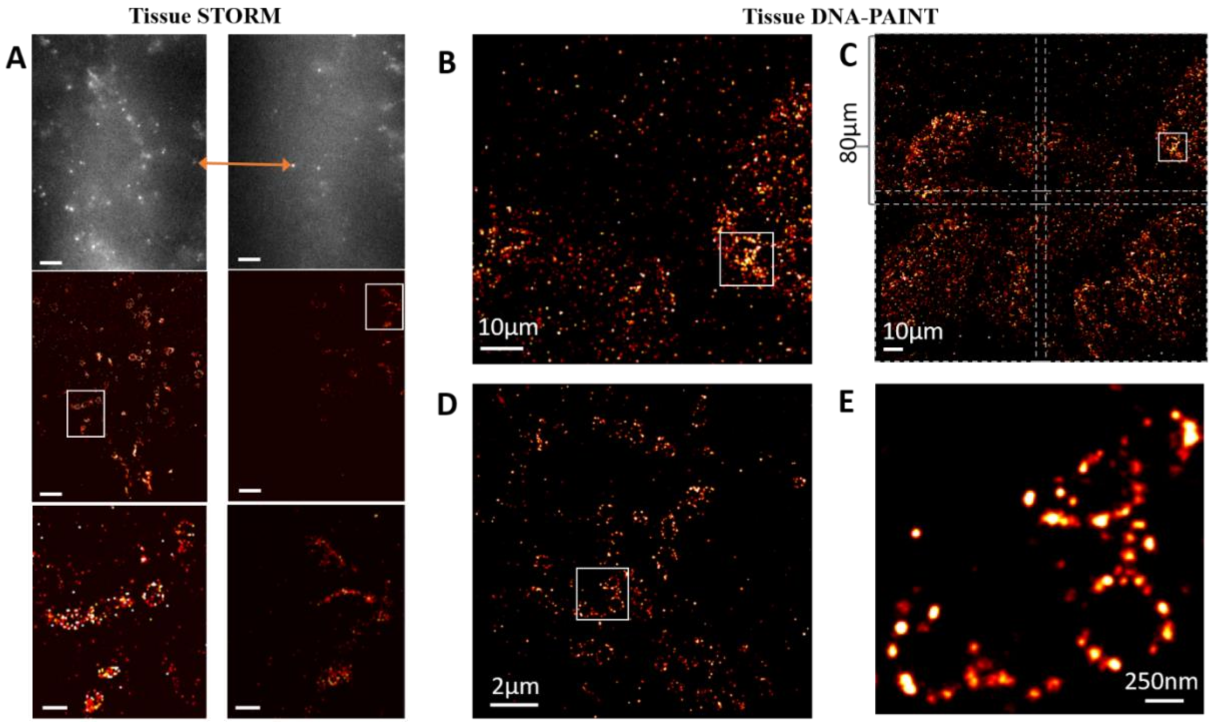


**Fig. S9.**

Representative images of both tissue STORM and tissue DNA-PAINT as acquired using a traditional objective-type TIRF microscope. (A) Single raw frame of tissue STORM and reconstructed view of Tom20-Alexa Flour 647 from pancreatic tissue FFPE for both an initially imaged FOV (left) and partially photobleached adjacent FOV (right). (B) Tissue DNA-PAINT of Tom20 within HER2+ breast cancer FFPE using largest FOV from our objective-type TIRF microscope of 80 µm x 80 µm. (C) Stitched 2x2 view of tumor boundary shows imaging of adjacent regions with tissue DNA-PAINT. (D) Magnified view from (B) of two cells within the tumor boundary. (E) Highest zoom-in showing rough mitochondria with poorly resolved boundaries. Tissue STORM images in (A) were acquired for 50,000 frames at 20ms exposure. Tissue DNA PAINT images in (D-E) were acquired for 50,000 frames at 100 ms exposure using 1 nm IS2-ATTO643 and 12.5 % EC. Scale bars in (A) are 5 µm, 5 µm and 1 µm for top, middle, and bottom rows respectively. Scale bar for (B) is 10 µm, (C) is 10 µm, (D) is 2 µm, and (E) is 250 nm.

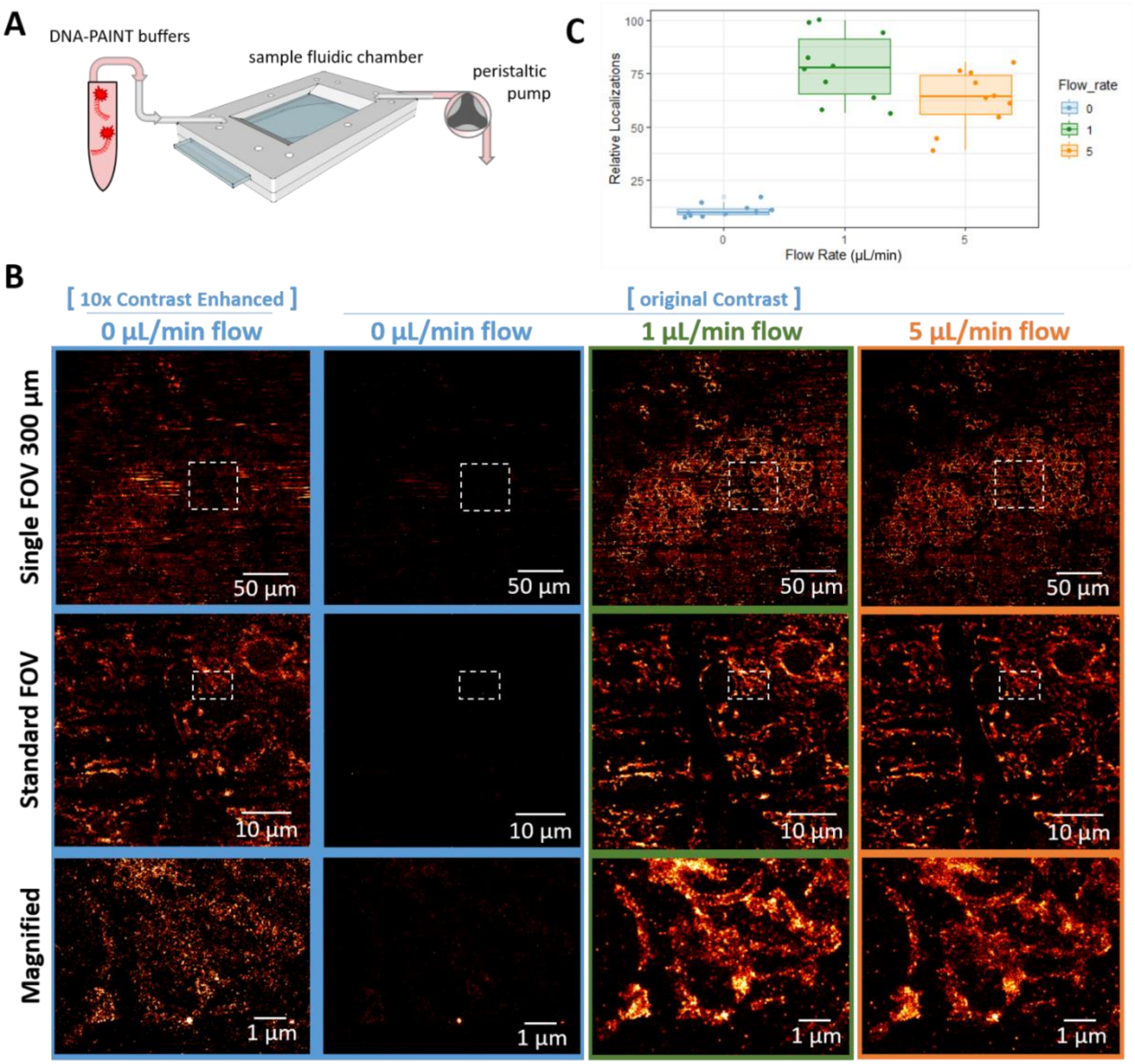


**Fig. S10.**

(A) Schematic overview of "microfluidics-enhanced DNA-PAINT" where a peristaltic pump is used to provide gentle flow within the thin (~35 µm) sample chamber. (B) Representative single FOVs of tissue DNA-PAINT imaged with or without "microfluidics enhanced DNA-PAINT" (top row). Zoom-in view from the single 300 µm FOV at a more standard FOV of 50 µm (middle row). Highest magnified view showing tissue mitochondria within a single cell (bottom row). Given the incredibly low relative signal without flow, we additionally showed a 10x contrast-adjusted views of the condition for each magnification (left column). Except for varying imaging buffer flow-rates, all images were acquired for 30,000 frames at 60 ms exposure using 500 pM IS2-ATTO643 and 7 % EC. Scale bars in (B) are 50 µm (top row), 10 µm (middle row), and 1 µm (bottom row). (C) Relative localization density for Tom20 from 10 FOVs imaged within a pancreatic FFPE tissue sample using the indicated flow rates 0, 1, and 5 µL/min.

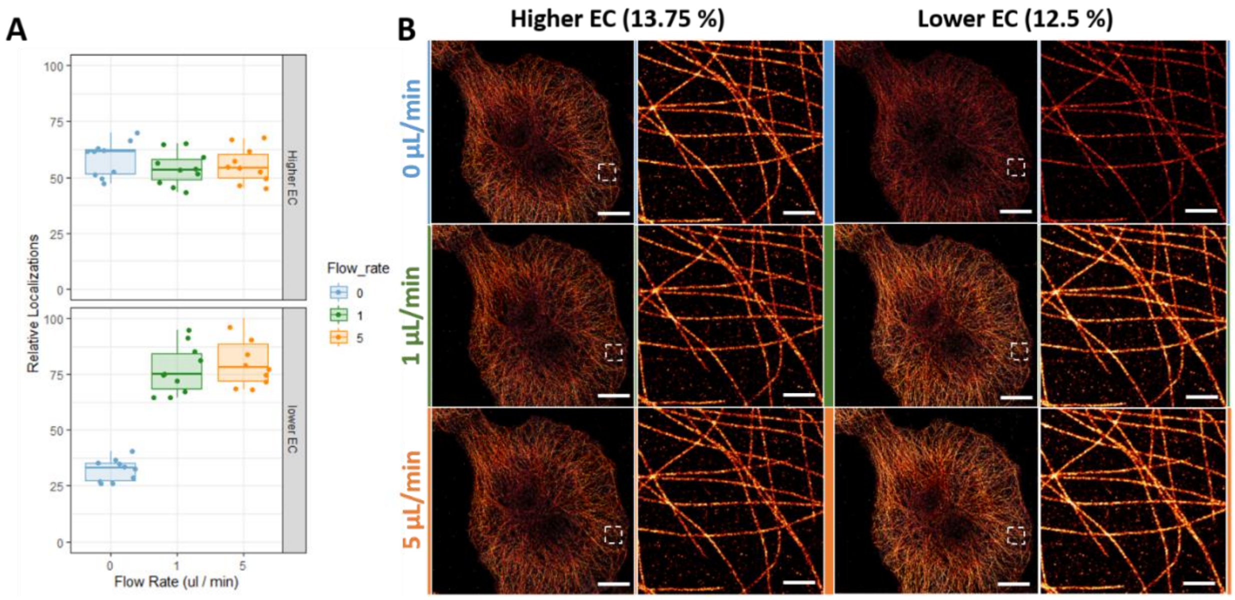


**Fig. S11.**

The relative effect of flow on cell-based imaging. (A) Relative localization density for microtubules from 10 FOVs imaged within a single Cos7 cell sample using the indicated flow rates of 0, 1, and 5 µL/min and either higher (13.75 %) or lower (12.5 %) EC. (B) Reconstructed views of cells imaged with or without “microfluidics enhanced DNA-PAINT”, with all images rendered with the same relative intensity. Except for varying imaging buffer % EC and flow-rates, all images were acquired for 20,000 frames at 30 ms exposure using 1nM IS1-ATTO643. Scale Bars in (B) are 5 µm and 500 nm.

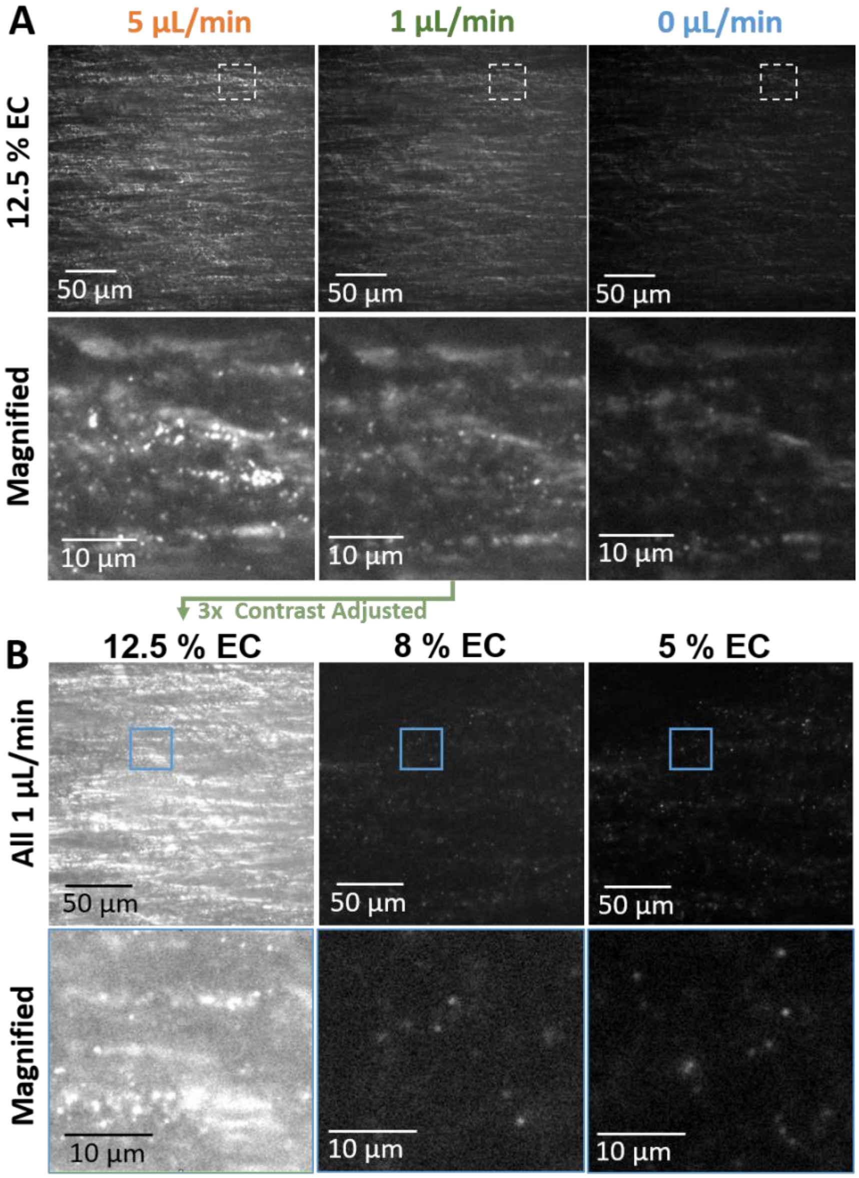


**Fig. S12.**

Representative raw frames of tissue DNA-PAINT on our PRIME-PAINT setup under varying EC concentration and "microfluidics enhanced DNA-PAINT" flow rates. (A) Attempt at using buffers optimized previously for cell imaging using PRIME-PAINT results in substantial diffusive background, exacerbated under increased flow rate during imaging. Magnified views show DNA-PAINT localizations, however high diffusive background lowers image quality. (B) Matching FOV images under 1 µL/min flow while varying EC concentrations (12.5 %, 8 %, & 5 %EC). Magnified views show a dramatic lowering of background at reduced EC, while DNA-PAINT localizations are more easily resolved. All frames within (A) or (B) are shown at the same relative intensity range, with the 12.5 % EC and 1 µL/min condition being 3x contrast-adjusted between (A) and (B). Raw frames shown were all acquired using 500 pM IS2-ATTO643 and a 60 ms exposure. Scale bars in (A, B) are 50 µm (top row) and 10 µm (bottom row).

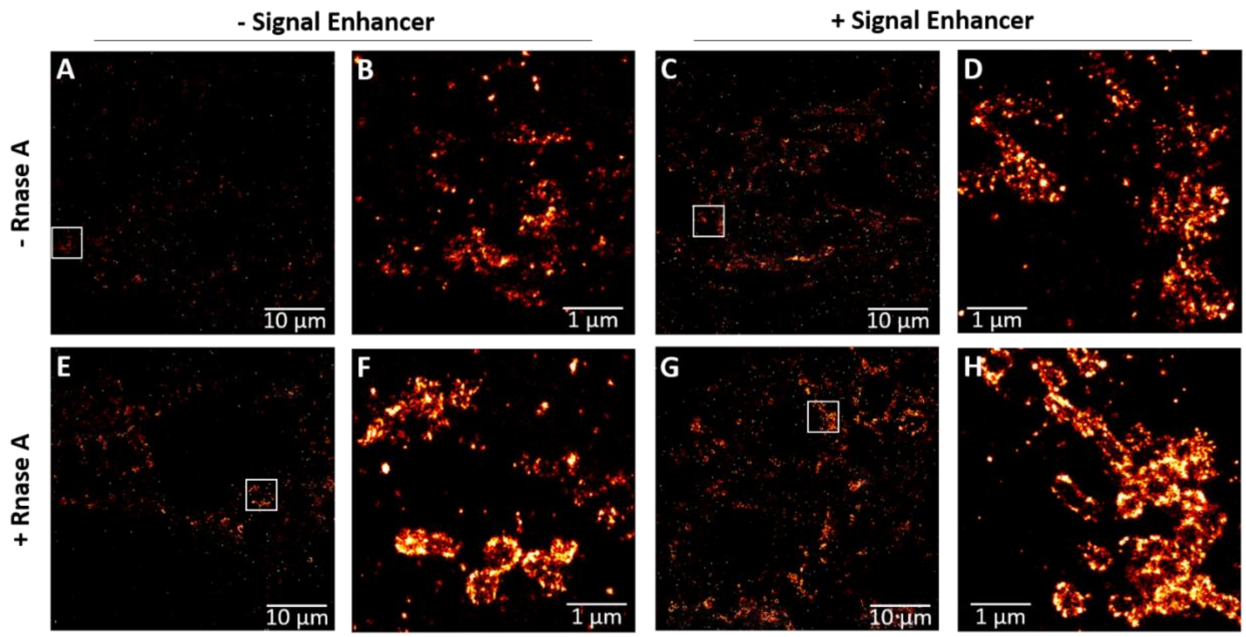


**Fig. S13.**

The effect of both RNase A and Signal Enhancer (SE) treatments on Tom20 labeling/imaging using “microfluidics enhanced DNA-PAINT” on pancreatic FFPE tissues. (A,C,E,G) Representative FOVs and (B,D,F,H) magnified regions from: control sample prepared identically to tissue STORM (A,B), sample with SE treatment prior to antibody labeling (C,D), sample with overnight RNase A treatment at RT prior to antibody labeling (E,F) and sample with sequential SE and RNase A treatment prior to antibody labeling (G,H). Outside of indicated treatment variations, all samples were prepared the same way and images were acquired using 30,000 frames at 60 ms exposure using 500 pM IS2-ATTO643 and 7 % EC. Scale bars are 10 µm in (A,C,E,G) and 1 µm in (B,D,F,H).

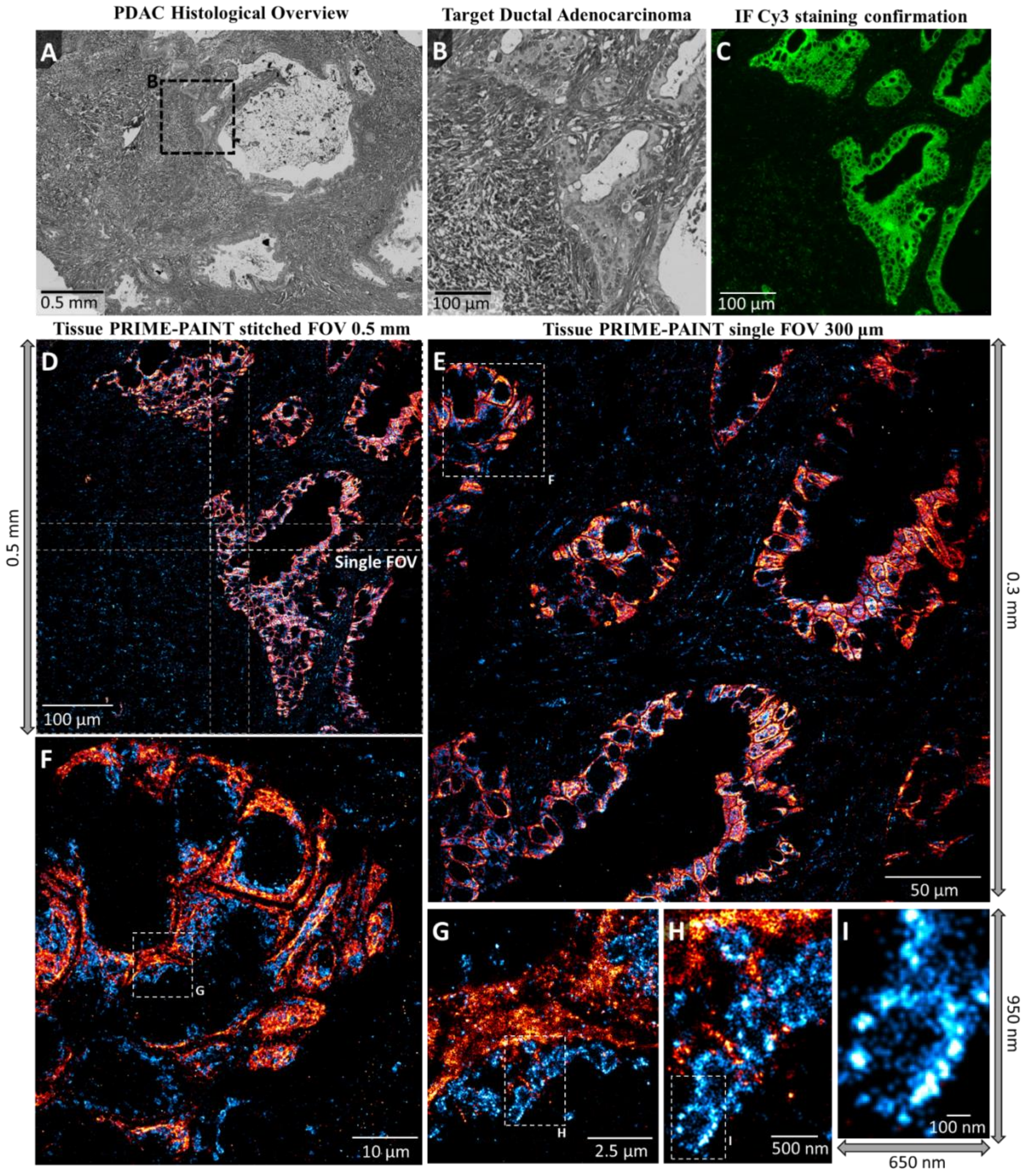


**Fig. S14.**

Additional FOV of tissue PRIME-PAINT on pancreatic cancer tissue sections. (A) Histological overview of moderately differentiated PDAC within desmoplastic stroma acquired at 20x magnification. (B) Targeted Ductal Adenocarcinoma for imaging with PRIME-PAINT. (C) Immunofluorescent confirmation of Cy3 signal from secondary antibodies conjugated to docking strand oligos showing strong pan-cytokeratin staining along the tumor and diffuse mitochondrial labeling within the tumor and adjacent stroma. (D) Stitched tissue PRIME-PAINT image of entire 0.5 µm wide ductal adenocarcinoma with both prognostic Pan-cytokeratin in red and mitochondrial Tomm20 in blue. (E) Single tissue PRIME-PAINT image obtained under a mild flow of imaging buffer (i.e., 'microfluidics-enhanced'). (F-I) Select serially magnified regions from (E) highlighting the increasingly fine features seen across different length scales. The entire tissue 2-target image (D-I) was acquired in 4 hours total (30,000 frames at 60 ms exposure for each target) at 1 µL/min flow and 500 pM IS1-ATTO643 and 7 % EC, and 500 pM IS2-ATTO643 and 7 % EC for pan-cytokeratin and Tom20 respectively. Scale bars are 500 µm in (A), 100 uµ in (B-D), 50 µm in (E), 10 µm in (F), 2.5 µm in (G), 500 nm in (H), and 50 nm in (I).
